# The vaccinia-based Sementis Copenhagen Vector COVID-19 vaccine induces broad and durable cellular and humoral immune responses

**DOI:** 10.1101/2021.09.06.459206

**Authors:** Preethi Eldi, Tamara H. Cooper, Natalie A. Prow, Liang Liu, Gary K. Heinemann, Voueleng J. Zhang, Abigail D. Trinidad, Ruth Marian Guzman-Genuino, Peter Wulff, Leanne M. Hobbs, Kerrilyn R. Diener, John D. Hayball

**Author notes:** These authors contributed equally to this work (Tamara H. Cooper, Natalie A. Prow). Corresponding authors: Dr Preethi Eldi; Clinical and Health Sciences Unit, University of South Australia, GPO Box 2471, Adelaide, SA, 5001, Australia.; Prof John Hayball; Sementis Limited, 25 North Tce, Hackney, SA, 5069, Australia.

## Abstract

The ongoing COVID-19 pandemic perpetuated by SARS-CoV-2 variants, has highlighted the continued need for broadly protective vaccines that elicit robust and durable protection. Here, the vaccinia virus-based, replication-defective Sementis Copenhagen Vector (SCV) was used to develop a first-generation COVID-19 vaccine encoding the spike glycoprotein (SCV-S).

Vaccination of mice rapidly induced polyfunctional CD8 T cells with cytotoxic activity and robust Th1-biased, spike-specific neutralizing antibodies, which are significantly increased following a second vaccination, and contained neutralizing activity against the alpha and beta variants of concern. Longitudinal studies indicated neutralizing antibody activity was maintained up to 9 months post-vaccination in both young and aging mice, with durable immune memory evident even in the presence of pre-existing vector immunity. This immunogenicity profile suggests a potential to expand protection generated by current vaccines in a heterologous boost format, and presents a solid basis for second-generation SCV-based COVID-19 vaccine candidates incorporating additional SARS-CoV-2 immunogens.

## Introduction

Severe acute respiratory syndrome coronavirus 2 (SARS-CoV-2) is responsible for coronavirus disease 2019 (COVID-19). It was first detected in late 2019^1, 2^ in Wuhan (Hubei Province, PRC) and has since spread globally with approximately 219 million confirmed cases and 4.5 million deaths as of September 2021^3^. Infection fatality rates are disproportionately high in aged individuals and individuals with co-morbidities such as diabetes, cardiovascular diseases, obesity, and immunosuppression^4–8^. The recent rapid spike in cases during the second and third waves of infection in several countries has been attributed to emerging variants of concern (VOC) with reported increased transmissibility of infection and/or disease susceptibility and severity associated with changes across the viral genome, in particular the spike protein^9–14^. Although initial counter measures such as social distancing, use of masks, and travel restrictions have played a major role in suppressing the spread of infection, vaccines currently continue to be the most effective means to control severe illness and fatality rates, and will play a major role in ending the enormous humanitarian and socioeconomic impact of the SARS-CoV-2 pandemic.

Currently 7 vaccines have been granted Emergency Use Listing by the World Health organization (WHO), with a further 112 in clinical development and 185 in pre-clinical development using a wide array of technologies including nucleic acid based vaccines (mRNA and DNA) ^15, 16^, replicating and non-replicating viral vectored vaccines^17–21^, inactivated viruses^22, 23^, and protein subunit vaccines^24^ using a range of new and repurposed adjuvants. Most of the vaccines in development, including all currently approved vaccines, exclusively target the SARS-CoV-2 spike glycoprotein to produce neutralizing antibodies to mediate protection. As a consequence of the unprecedented speed by which frontrunner vaccines were developed, there remains knowledge gaps in vaccine-mediated protection such as duration of protection, efficacy in specific clinically vulnerable populations, rare side effects affecting safety, induction and relative importance of cell-mediated immunity, cross-protection against other coronaviruses, and control of viral shedding and transmission. In addition, vaccines must meet the needs for rapid and successful deployment within the context of population-scale vaccination programs, with specific considerations given to large scale manufacturability and distribution logistics. This highlights the need for continued COVID-19 vaccine development strategies using novel technologies that are safe and build upon the successes of first-generation vaccines to provide robust, durable and broadly protective efficacy against variants as well as continue to address key logistical challenges.

The Sementis Copenhagen Vector (SCV) is a rationally designed, replication-defective viral vector technology based on the Copenhagen strain of vaccinia virus (VACV). Uniquely for a VACV-based vector, it has also been paired with a proprietary manufacturing cell line (MCL) to generate a platform system that can facilitate large scale and facile manufacturability of all SCV-based vaccines^25^. The SCV was generated by the targeted deletion of an essential viral assembly gene (*D13L*)^26, 27^ to prevent viral replication whilst retaining the powerful immunogenicity and large payload capacity of the parental virus. With sites B7R/B8R, C3L, A39R and A41L designated as antigen insertion-sites, it is also an ideal vector technology for multi-antigen and multi-pathogen vaccines^28^. Systems vaccinology studies have demonstrated that SCV vaccination results in a localised Th1 signature and significant immunogen expression at the injection site^29^, with advanced preclinical studies showing a fully attenuated and safe vaccine capable of inducing robust and long-lived antigen-specific antibody and CD8 T cell responses equivalent to those elicited by replication competent VACV^25^. Vaccine efficacy of SCV vaccines has also been established in mouse models of disease and non-human primate immunogenicity and infectious disease challenge studies^25, 28, 30^.

In this study, a first-generation SCV vaccine encoding the full-length, native spike glycoprotein (SCV-S) was constructed, with cellular expression and cell surface anchorage of the spike protein in host cells confirmed. Detailed immunological analyses of vaccine-mediated immune responses demonstrated robust and long-lived spike-specific CD8 T cell and neutralizing antibody responses in young and aging mice, including inbred and outbred strains. Following a second booster dose, circulating neutralizing antibody levels were sustained without any discernible decay over a nine-month period. Assessment of long-lived CD8 T cell and antibody-secreting cell compartments confirmed induction of durable spike-specific immune memory following vaccination. Therefore, this study presents evidence that SCV-based COVID-19 vaccines can mediate robust and durable serological and cellular priming which supports progression towards efficacy studies of second-generation vaccine candidates incorporating additional SARS-CoV-2 antigens, to build synergistic layers of additional protection.

## Results

### Construction and characterization of a SCV expressing the SARS-CoV-2 spike protein

The proprietary SCV vaccine platform technology comprises (1), a viral vector that is unable to produce infectious viral progeny through targeted deletion of an essential viral assembly protein (D13) from VACV, which maintains amplification of the viral genome and late gene expression of introduced vaccine antigens for immune stimulation. This is combined with (2), a manufacturing cell line based on Chinese hamster ovary (CHO) cells that constitutively express D13, for virion assembly, and CP77, an essential VACV host-range protein for CHO cells that provides replication capability for SCV vaccine production in suspension cultures using established commercial manufacturing technologies and facilities (Figure 1a).

**Figure 1:**
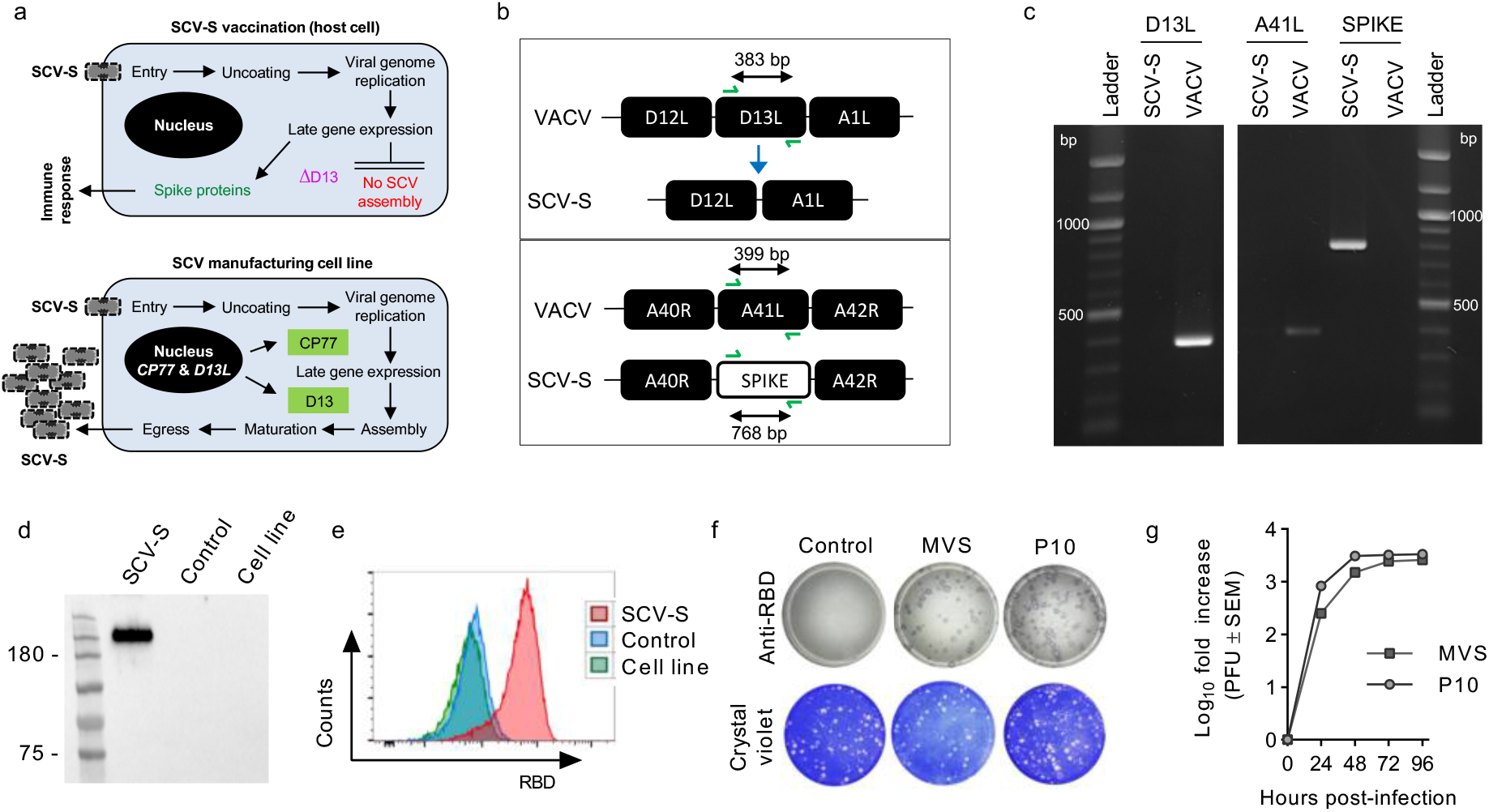
Vaccine construction and *in vitro* characterization. (**a**) Targeted deletion of *D13L* in SCV-S renders the viral vector unable to generate viral progeny by preventing virion assembly but allows for amplification of the SCV-S genome and late gene expression for production of spike protein to drive the vaccine-specific immune response (top panel). *In trans* provision of D13 and expression of the host range protein CP77 in the CHO-based manufacturing cell line allows for cell infection and rescue of virion assembly, allowing production of progeny for vaccine manufacture (bottom panel). (**b**) Schematic representation of SCV-S construction showing the *D13L* deletion and *A41L* substitution sites, with site specific primers and the expected band size indicated. (**c**) Viral DNA from VACV and SCV-S was used for PCR analysis using the site specific primers shown in (b) to confirm the deletion of D13L and insertion of spike. (**d**) Immunoblot analysis of spike antigen expression in SCV-S and control vector infected 143B cells using an anti-S1 antibody. (**e**) Surface analysis of RBD expression in SCV-S or control vector infected or uninfected MCL cells by flow cytometry using an anti-RBD antibody. (**f**) Immunostaining for RBD expression in STO1-33 cell monolayers infected with control vector, master viral seed (MVS) or passage 10 (P10) stocks of SCV-S. Crystal violet was used to visualize plaques. (**g**) Manufacturing cell line was infected with MVS and P10 stocks of SCV-S vaccine at an MOI of 0.01 PFU, and viral titers were determined by plaque assay at the indicated times (n=3 replicates).

To generate a spike encoding SCV vaccine (SCV-S), the full-length SARS-CoV-2 spike gene from the original Wuhan isolate-1 was placed under the control of a synthetic VACV early/late promoter^31^ and introduced by homologous recombination into the A41L deletion locus of the SCV (Figure 1b). PCR analysis of recombinant viral DNA confirmed site-specific insertion of the heterologous SARS-CoV-2 spike sequence into SCV and absence of *D13L* (Figure 1c). Authentic expression of the spike protein was confirmed by infecting non-permissive human 143B cells^25^ with SCV-S or parent control vector (SCV containing no vaccine antigen) and detection of S1 production by immunoblot analysis (Figure 1d), as well as surface expression of the receptor binding domain (RBD) in infected cells by flow cytometric analysis (Figure 1e).

The conversion of research grade vaccine stock to cGMP-compliant master and working virus seed (MVS and WVS) stocks requires several passages, followed by subsequent expansion during manufacturing for large scale SCV vaccine production. Therefore, the genetic stability of SCV-S was evaluated by serial passaging of the MVS up to 10 times (P10), which extends at least five-fold beyond the expected expansion requirement to reach commercial-scale vaccine batch production. Immunostaining of MVS and P10 infected cell monolayers using anti-RBD antibody confirmed spike protein expression in all foci of viral infection (Figure 1f), with similar viral titer amplification ratios observed up to 72 hrs post-infection (Figure 1g). Genetic integrity and sequence authenticity was also confirmed by next-generation sequencing of MVS and P10 stocks (data not shown), which together with characterised expression indicated the transgene stability and genetic integrity of the SCV-S vaccine candidate.

### SCV-S vaccination stimulates rapid and functional cell-mediated immunity

The capacity of SCV-S to induce an early spike-specific cellular immune response was evaluated in C57BL/6J mice. One week after vaccination, recovered splenocytes were stimulated with overlapping peptide pools spanning the S1 and S2 subunits of the SARS-CoV-2 spike protein and antigen-specific T cell responses evaluated by IFN-γ ELISPOT. Vaccination with SCV-S induced significant S1- and S2-specific IFN-γ spot forming units (SFU) compared to mice vaccinated with control vector (Figure 2a). Consistent with the ELISPOT results, significantly elevated populations of S1- and S2-specific IFN-γ producing CD8+ T cells were detected in SCV-S vaccinated mice compared to controls by intracellular cytokine staining (Figure 2b).

**Figure 2:**
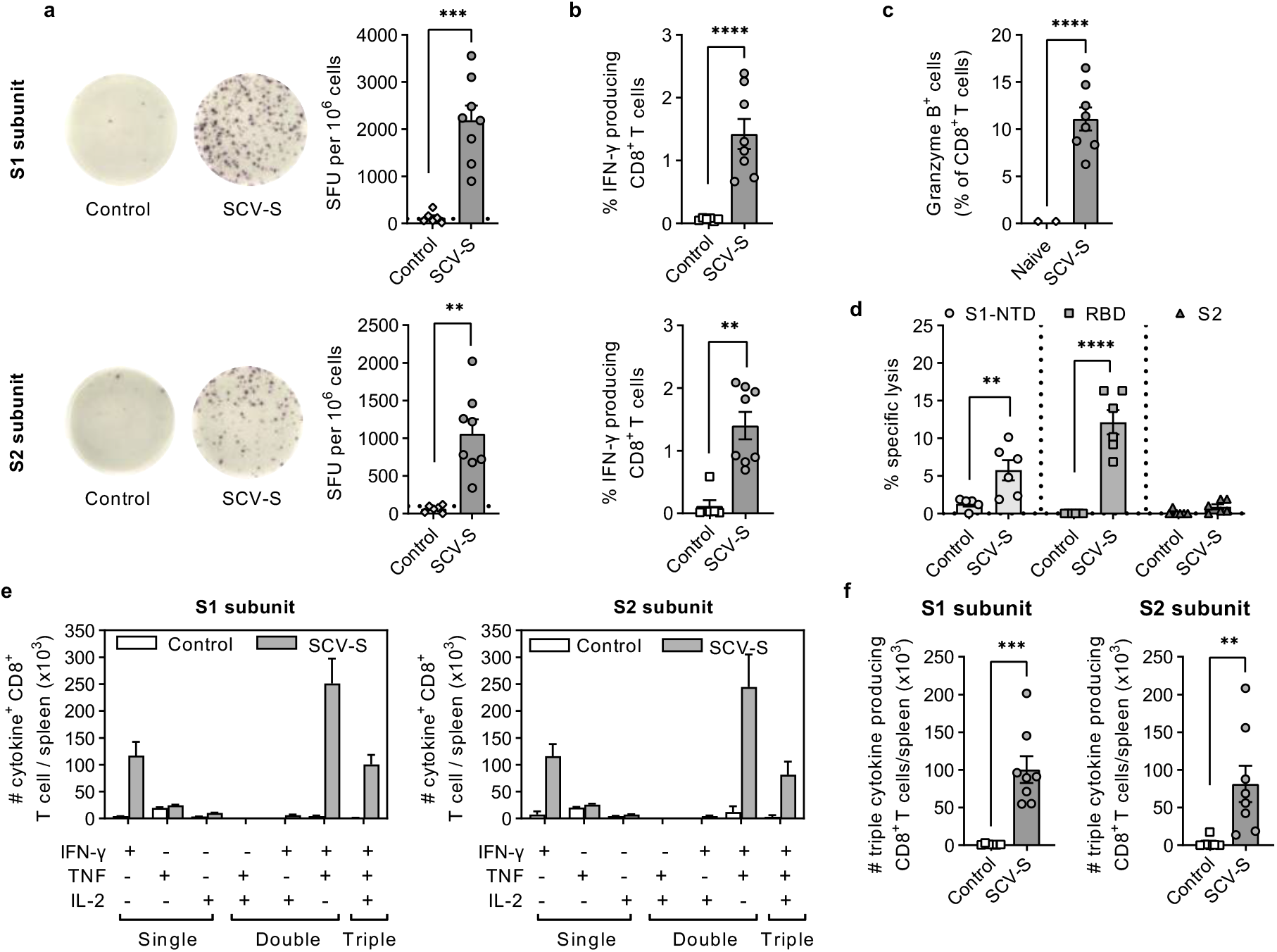
Early CD8 T cell responses following SCV-S vaccination. (**a**) Groups of C57BL/6J female mice (n=5) were vaccinated with 10^7^ PFU of SCV-S or control vector and CD8 T cell responses assessed 1 week later. S1 and S2-specific IFN-γ spot-forming units (SFU) quantitated by ELISPOT after stimulation of splenoctyes with peptide pools (15AA length with 11mer overlaps) spanning the S1 and S2 subunits of the spike protein. (**b**) Frequency of IFN-γ^+^ CD8^+^ T cells specific for S1 and S2 subunit enumerated by intracellular cytokine staining in cohorts described in (a). (**c**) Frequency of intracellular Granzyme B^+^ CD8^+^ T cells in C57BL/6J mice (equal gender; n=8) vaccinated with SCV-S for 7 days or naïve controls. (**d**) Percent specific lysis of MC57G target cells pulsed with S1-NTD-, RBD-, and S2-subunit specific peptides assessed by standard ^51^Cr release *ex vivo* CTL assay 7 days after vaccination of C57BL/6J mice (n=6) with SCV-S or control vector. **(e)** Absolute number of single, double, and triple cytokine producing CD8^+^ T cells in S1 and S2 stimulated splenocytes from experiment in (a), with **(f)** S1- and S2-specific triple cytokine producing CD8 T cells presented against that found in control vector groups. Symbols represent individual mice and bars show the mean ± SEM. Data was log transformed and unpaired *t*-test with Welch’s correction was used for statistical analysis. **p<0.01; ***p<0.001; ****p<0.0001

Effector CD8 T cells with cytotoxic potential produce granzyme B, a serine protease that is capable of mediating target cell lysis^32^, and a significant population of CD8^+^ T cells were shown to express granzyme B in response to SCV-S vaccination (Figure 2c). To confirm the SCV-S vaccine induced a functional cytotoxic T lymphocyte response, day 7 splenocytes were incubated with radiolabelled target cells pulsed with either S1 N-terminal domain (S1-NTD)-, RBD-, or S2-specific spike subunit peptides (Supplementary Table 1). Significant cytolytic activity was detected against S1-NTD and RBD target cells (Figure 2d). The absence of functional activity against the S2 subunit suggested that the SCV-S vaccine primarily triggers a S1-specific cytotoxic T cell profile. Together these results indicated that the SCV-S vaccine induced an early expansion of spike-specific effector CD8 T cells that display the potential to reduce the viral burden before the humoral arm of the immune response can be well established.

Previous studies have demonstrated CD8 T cells that produce multiple cytokines have enhanced effector functions and support maturation of an antigen-specific memory T cell population^33, 34^. Therefore, S1- and S2-specific cytokine producing CD8^+^ T cells were profiled into seven distinct populations based on the production of IFN-γ, TNF, IL-2, and their combinations (Supplementary Figure 1). Polyfunctional IFN-γ^+^ TNF^+^ cells dominated the responding S1- and S2-specific T cell population, followed by IFN-γ^+^ TNF^+^ IL-2^+^ (triple cytokine producing) and single IFN-γ^+^ producing cell population (Figure 2e). Furthermore, significant numbers of S1- and S2-specific triple cytokine producing CD8^+^ T cells (Figure 2f), produced more cytokines on a per cell basis compared to double and single cytokine producing CD8^+^ T cells (Supplementary Figure 2).

### SCV-S vaccination induces spike-specific binding and neutralizing antibody responses

Immunogenicity of SCV-S was evaluated in C57BL/6J mice, with kinetics of spike binding antibody responses assessed up to day 28 post-vaccination. S1 and S2 IgM binding titers peaked at day 7 post-vaccination then returned to baseline levels by day 20 (Figure 3a; top panels). Class-switched S1 IgG binding titers were detected in all mice by day 9 post-vaccination, with the appearance of S2 IgG binding antibodies delayed to days 15-19 post-vaccination (Figure 3a, lower panels). The World Health Organisation (WHO) target product profile for COVID-19 vaccines^35^ and current understandings in the field indicate that a Th1-biased humoral and cellular immune responses is required to prevent vaccine-associated enhanced respiratory disease^36–38^. Therefore, subclass profiling of S1 and S2 binding antibodies was assessed in day 28 post-vaccination serum samples. A strong IgG2c response with an associated IgG2c/IgG1 ratio of > 1 was observed confirming a Th1-biased spike-specific antibody response (Figure 3b and Supplementary Figure 3).

**Figure 3:**
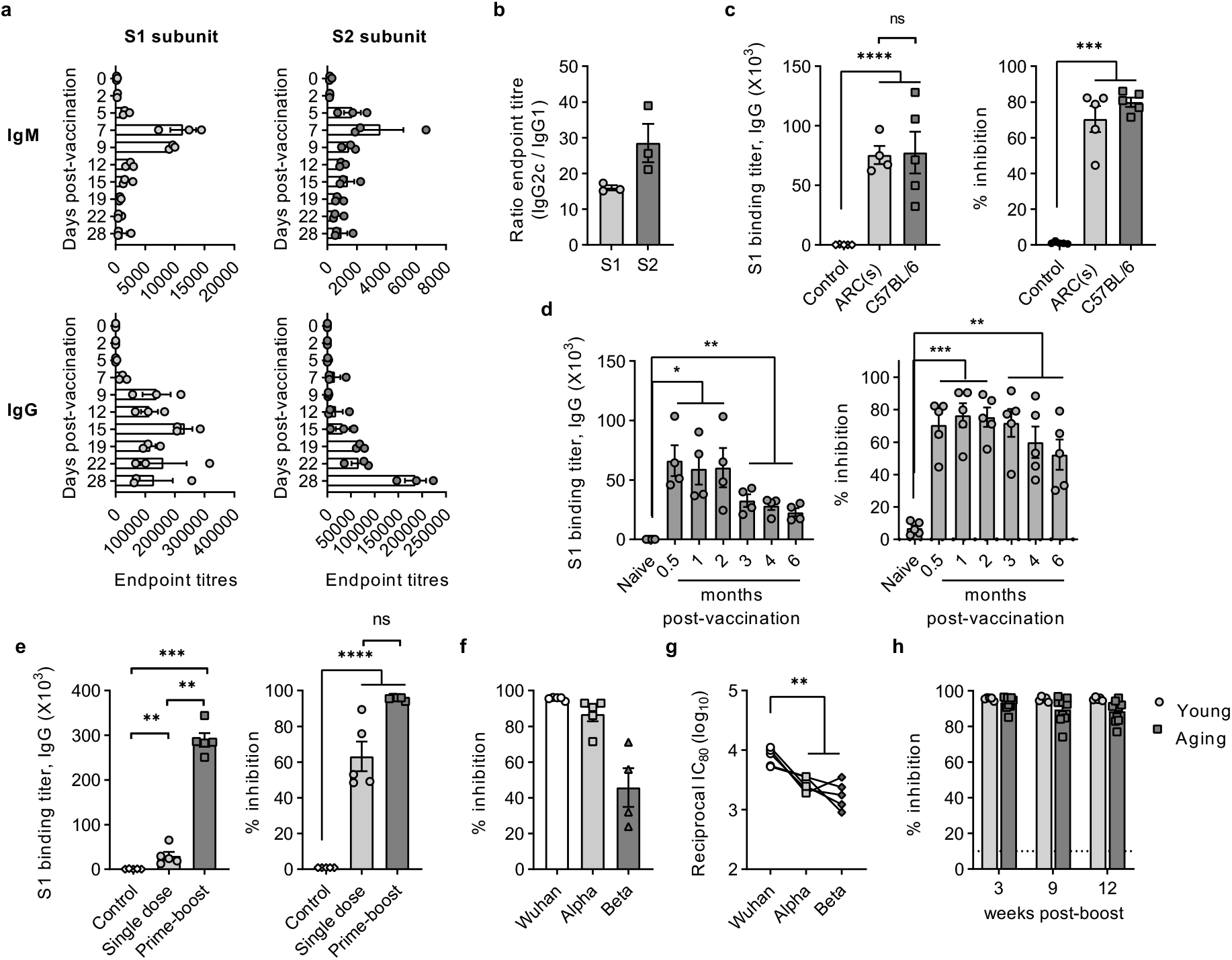
Spike-specific antibody and SARS-CoV-2 neutralization responses following SCV-S vaccination. (**a**) S1 and S2 subunit endpoint IgM and IgG ELISA titers determined from serum of female C57BL/6J mice (n=3) at the indicated times after a single vaccination with 10^7^ PFU of SCV-S. (**b**) Ratio of S1- and S2-specific IgG2c to IgG1 endpoint ELISA titers determined 28 days post-vaccination. (**c**) S1 IgG ELISA binding titers (left panel) and cPass^TM^ neutralization titers (right panel) in outbred ARC(s) and inbred C57BL/6J female mice (n=5) 21 days after a single vaccination with 10^7^ PFU of SCV-S or vector control. (**d**) S1-specific endpoint IgG ELISA titers (left panel) and neutralization titers (right panel) in female outbred ARC(s) mice (n=5) at the indicated times after a single vaccination with 10^7^ PFU of SCV-S or vector control. (**e**) S1-specific IgG ELISA (left panel) and neutralization titers (right panel), 50 days after single dose (day 0) or prime-boost (day 0 and 28) vaccination of female C57BL/6J mice (n=5) with 10^7^ PFU SCV-S or control vector. (**f**) neutralizing activity against lenti-SARS-CoV-2-S pseudoviruses bearing spike protein from either the Wuhan reference strain, the Alpha, or the Beta variant, and (**g**) corresponding IC80 titers, shown for prime-boost samples. (**h**) Neutralization titers in young (6-8 week old) and aging (9-10 month old) C57BL/6J mice at the indicated times after prime-boost vaccination with 10^7^ PFU SCV-S. Symbols represent individual mice and bars show the mean ± SEM. Data was log transformed and statistical significance determined using Brown-Forsythe and Welch ANOVA with Dunnett T3 multiple comparison test. *p<0.05; **p<0.01; ****p<0.0001

Outbred strains of mice are more representative of the genetic variability in the general human population and therefore the dominant S1 binding antibody responses in inbred and outbred strains of mice were compared at day 21 post-vaccination. S1 IgG binding titers were comparable between the two strains of mice, were significantly higher than the controls, and translated into significant levels of antibody-mediated neutralization capacity (Figure 3c).

To understand the long-term kinetics of the spike-specific antibody response, serum samples from outbred ARC(s) mice were evaluated up to 6 months post-vaccination. S1 binding titers remained significantly elevated compared to naïve mice at all time-points tested, although a trend in reduced titer was observed from 3 months. Significant neutralizing activity was also maintained, albeit with a small steady decline after 3 months (Figure 3d). A similar trend was observed in C57BL/6J mice, with significant S1-specific antibody binding titers and neutralizing activity maintained up to 12 weeks post-vaccination, although neutralization activity had halved from 3 week levels by 12 weeks (Supplementary Figure 4). Therefore, to stabilise the waning antibody responses, a 4-week homologous prime/boost vaccination strategy was assessed in C57BL/6J mice. At 3 weeks post-boost, a significant increase in both S1 IgG binding titers and neutralizing activity was detected in the prime-boost cohort compared to single dose and control vector vaccinated mice (Figure 3e). A positive correlation between the S1-specific IgG binding titers and neutralization activity was confirmed (Supplementary Figure 5).

Given the emergence of SARS-CoV-2 variants that can exhibit increased replicative and transmission fitness, the SCV-S vaccine-generated neutralization antibody response was assessed against the recognised Alpha and Beta VOCs. First, the capacity to block the interaction between viral RBD and the host angiotensin-converting enzyme 2 (ACE2) receptor was assayed using the RBD protein from the original Wuhan Isolate-1, N501Y Alpha VOC or the E484K, K417N and N501Y Beta VOC. All serum samples inhibited the binding between the ACE2 receptor and the RBD from original Wuhan isolate-1, with a 1.1- and 2.1-fold reduction in inhibition detected for the Alpha and Beta VOCs (Figure 3f). Next, using SARS-CoV-2 pseudotyped lentiviruses containing the spike sequence of either the Wuhan isolate-1, Alpha, or Beta VOCs, the effect of all the mutations in the spike protein on the serum neutralizing capacity was examined. A similar result, with a significant 3.2- and 4.1-fold reduction in the IC_80_ (80% inhibitory concentration) was noted for the Alpha and Beta VOCs compared to the original Wuhan isolate-1 (Figure 3g).

The persistence of neutralizing antibody responses against SARS-CoV-2 is important in all populations, including the aging population. Therefore, the durability of the SCV-S induced neutralizing antibody activity was evaluated in both young (6-8 weeks old) and aging (9-10 months old) mice. Functional neutralizing antibodies were maintained without any significant decrease in titer up to the termination of the study at three months post-boost vaccination (a total of 4 months from first vaccination dose). Importantly, no significant differences were detected in the neutralizing capacity between the young and aging mice (Figure 3h). In summary, these results indicate that administration of SCV-S in a prime-boost vaccination strategy induces a robust and durable SARS-CoV-2 spike-specific antibody response with significant neutralizing capacity against the original Wuhan isolate-1, and both Alpha and Beta VOCs.

### Prime-boost vaccination with SCV-S enhances antigen-specific memory T cell responses

To examine SCV-S-induced cellular immune responses, C57BL/6J mice were immunised in a single dose (day 0) or prime-boost (day 0 and 28) vaccination protocol, with spike-specific memory CD8^+^ T cell populations studied three months later. Mice vaccinated with control vector in a prime-boost strategy were used as appropriate controls. Following a single dose of the vaccine, S1-NTD-specific IFN-γ^+^ SFU were detected above the background determined by control vector vaccinated mice, however a significant 4.3-fold increase over controls was noted with a prime-boost regimen (Figure 4a). This correlated with an increased frequency of IFN-γ^+^ producing CD8^+^ T cells and absolute numbers of polyfunctional IFN-γ^+^ TNF ^+^ IL-2^+^-expressing CD8^+^ T cells, with the prime-boost regimen significantly elevating the response over that detected in single dose and control vaccinated mice (Figure 4b). A similar significant increase in RBD-specific and S2-specific IFN-γ^+^ SFU, and IFN-γ-expressing and polyfunctional CD8^+^ T cells was noted in the prime-boost vaccinated mice compared to the single dose and control vaccinated mice; however a single dose also induced a significant population of RBD-specific and S2-specific IFN-γ^+^ CD8^+^ T cells and triple cytokine producing cells (Figures 4c-4f).

**Figure 4:**
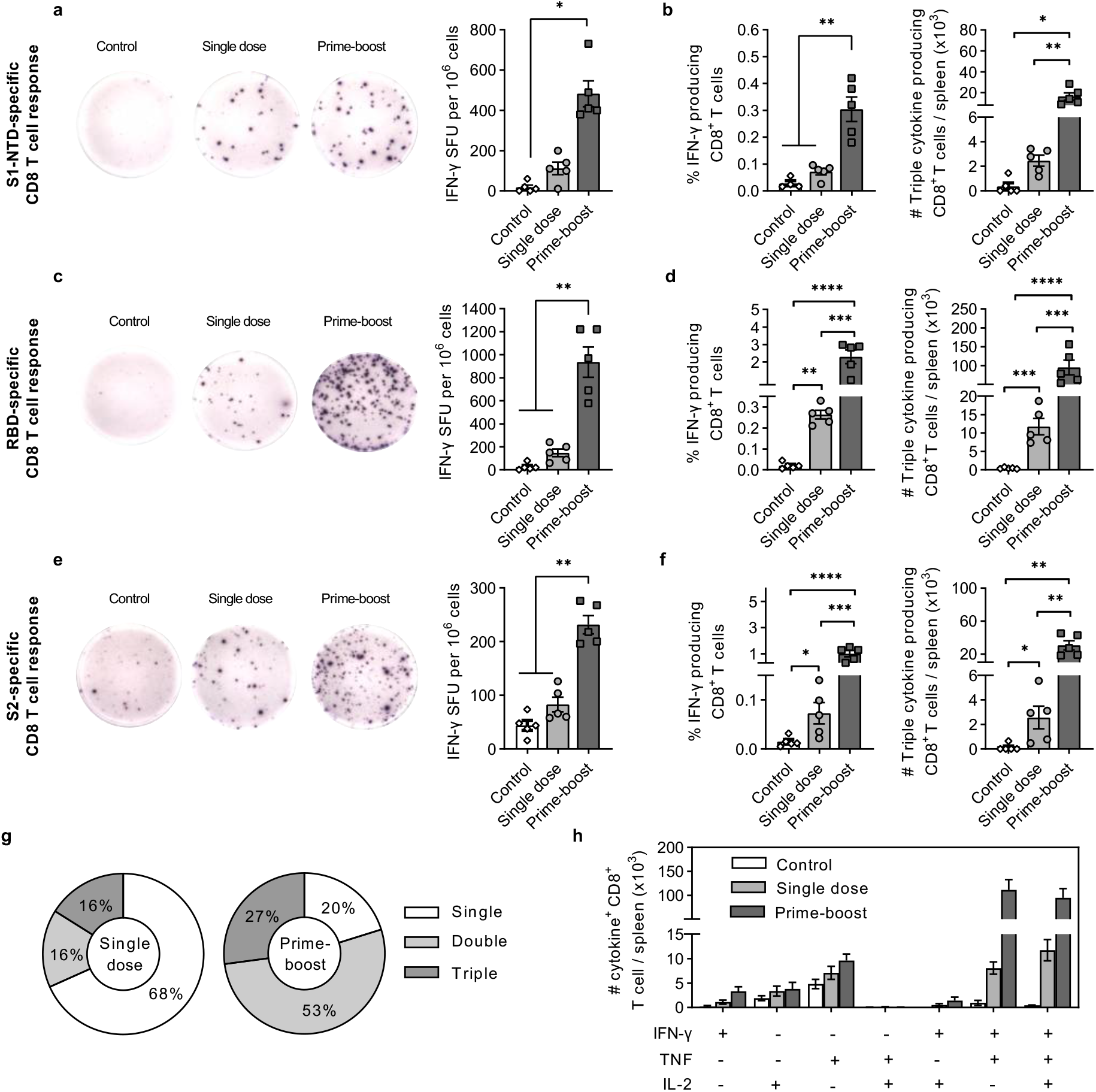
Spike-specific CD8 T cell responses 3 months post SCV-S vaccination. Groups of female C57BL/6J mice (n=5) were vaccinated in a single dose (day 0) or prime-boost strategy (day 0 and 28) with SCV-S. Mice vaccinated with control vector on day 0 and 28 were used as controls. At day 120, splenocytes were stimulated with peptide pools (15AA length with 11mer overlaps) spanning the spike protein to examine antigen-specific T cell responses. **(a,c,e)** IFN-γ SFU specific forS1-NTD, RBD-, and S2-specific subunit regions of the spike protein was quantitated by ELISPOT. **(b,d,f)** Corresponding frequency of IFN-γ^+^ CD8^+^ T cells and absolute numbers of triple cytokine (IFN-γ^+^ TNF ^+^ IL-2^+^) producing CD8 T cells as enumerated by intracellular cytokine staining and FACS analysis. **(g)** Graphs showing the mean frequency and (**h**) absolute numbers of RBD-specific single (IFN-γ^+^, TNF ^+^ or IL-2^+^), double (TNF ^+^ IL-2^+^, IFN-γ^+^ IL-2^+^ or IFN-γ^+^ TNF ^+^) and triple cytokine (IFN-γ^+^ TNF^+^ IL-2^+^) cells within the cytokine positive CD8 T cell compartment. Symbols represent individual mice and bars show the mean ± SEM. Data was log transformed and statistical significance was determined using Brown-Forsythe and Welch ANOVA with Dunnett T3 multiple comparison test. *p<0.05; **p<0.01; ***p<0.001; ****p<0.0001

Analysis of the total spike-specific CD8^+^ T cell IFN-γ^+^ responses revealed an immunodominance of RBD-specific responses, with ∼65% of the total spike-specific responses targeting the RBD region of the spike protein. Further cytokine profiling of this RBD-specific CD8^+^ T cell population revealed that the prime-boost vaccination regimen significantly increased the mean frequency of multifunctional T cells from 32% with a single dose, to 80% with a prime-boost. This consisted of an average 3.3-fold increase in double positive, and 1.7-fold increase in triple cytokine positive cells compared to single dose vaccinated mice (Figure 4g and 4h).

Vaccine antigen-specific cellular immune responses were also examined in aging mice vaccinated once or in a prime-boost strategy. Similar trends to that seen in young mice was observed, with prime-boost vaccination inducing significantly higher S1-NTD-, RBD-, and S2- specific IFN-γ producing cells compared to single dose and control vaccinated mice.

Additionally, the proportions of single TNF-secreting CD8^+^ T cells and numbers of triple cytokine producing CD8^+^ T cells were significantly increased over that induced by a single dose, as was the mean frequency and absolute number of multifunctional T cell proportions (Supplementary Figure 6).

Vector-specific memory T cell responses were also assessed, and as expected, prime-boost SCV-S and control vector vaccinated mice had significantly higher levels of vector-specific IFN-γ SFU, IFN-γ^+^ CD8^+^ T cells, and triple cytokine positive cells compared to single dose vaccinated mice (Supplementary Figure 7). Overall, prime-boost vaccination significantly increased the CD8^+^ short lived effector T cell (T_SLE_) and long-lived effector memory T cell (T_EM_) populations in general. No significant changes were observed in the central memory T cell (T_CM_) compartment (Supplementary Figure 8). Altogether, these results confirm that prime-boost vaccination with SCV-S enhances the spike-specific memory CD8^+^ T cell response in both young and aging populations of mice.

### Pre-existing vector immunity constrains the magnitude of cellular immune responses, but has no impact on spike-specific humoral immune responses

The presence of VACV-specific memory immune responses, particularly in a small cohort of the aged population that had previously received a smallpox vaccination, may impact on the induction of vaccine antigen-specific cellular and humoral immune responses from recombinant SCV vaccines. Therefore, a mouse model of robust pre-existing vector immunity was established to study its impact on spike-specific immune responses following vaccination with SCV-S. Mice were administered a single dose of replicative VACV, with presence of vector-specific antibody responses confirmed 6 weeks later (Figure 5a). After eight weeks, mice were vaccinated with SCV-S vaccine in a 4-week prime-boost strategy, and the magnitude and quality of spike-specific antibody and T cell responses evaluated. Naïve mice were used as controls for vector-specific immune responses and mice vaccinated with control vector were used as controls for antigen-specific responses. Two weeks post SCV-S booster dose administration, no significant differences in S1- and S2-specific IgG binding titers between cohorts with (+) or without (-) pre-existing immunity were observed (Figure 5b). Consistent with the binding titers, SARS-CoV-2 neutralization activity was comparable between the two groups of mice (Figure 5c). Importantly, S1 IgG binding titers and neutralizing activity were not impacted by pre-existing immunity even at three months post SCV-S vaccination (Figure 5d).

**Figure 5:**
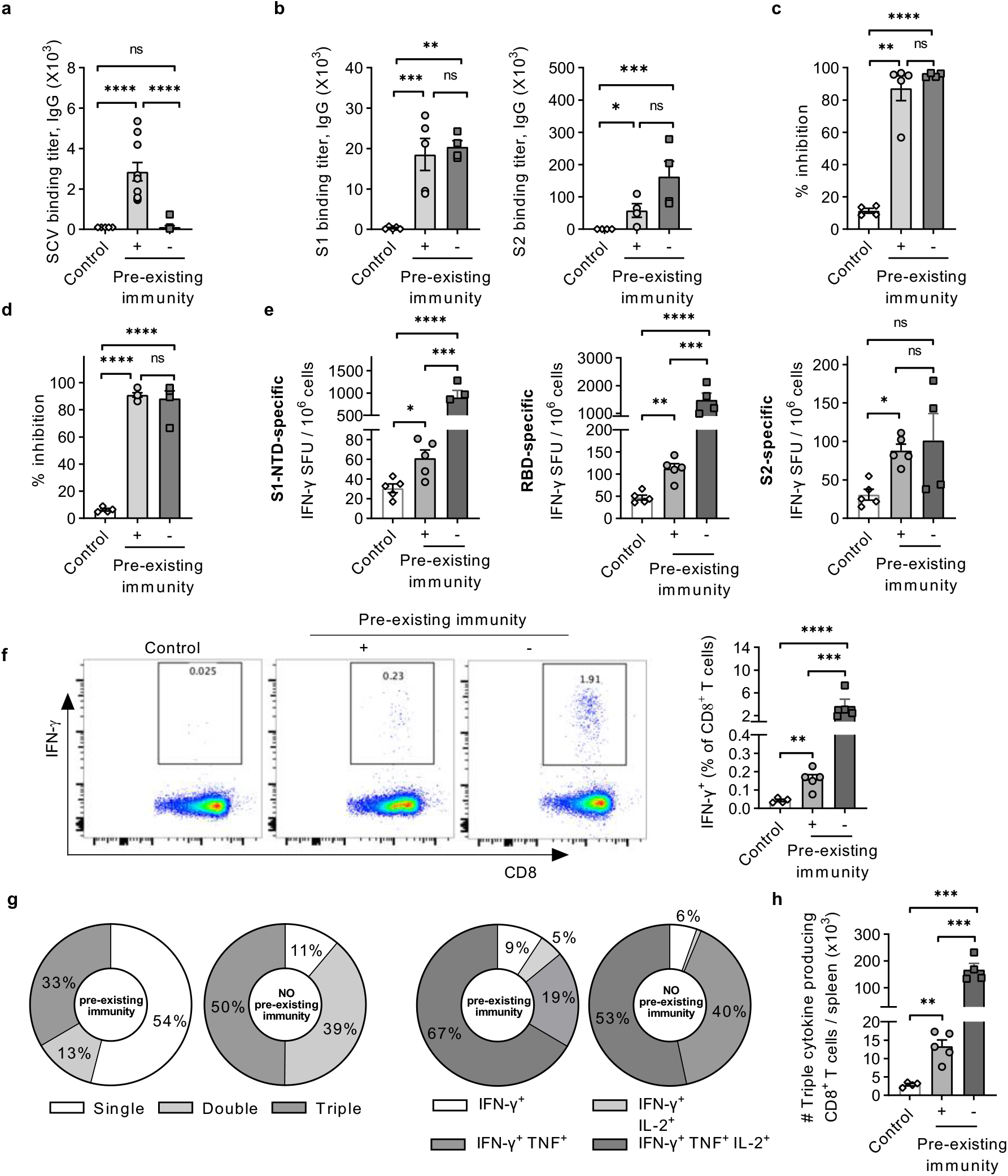
Impact of pre-existing vector immunity on spike-specific antibody and CD8 T cell responses. Cohorts of C57BL/6J mice containing both genders (n=5 in total) were vaccinated at day -60 with VACV at 10^5^ PFU or diluent only to generate groups with and without pre-existing immunity. Mice were subsequently vaccinated with 10^7^ PFU SCV-S at day 0 and 28. (**a**) Vector-specific endpoint IgG ELISA titers 28 days after exposure to VACV or diluent control. (**b**) S1 (right panel) and S2 (left panel) IgG ELISA, and (**c**) neutralization titers 50 days after vaccination. (**d**) Maintenance of neutralization activity 3 months post-vaccination. (**e**) S1-NTD-, RBD-, and S2- specific IFN-γ SFU were quantitated by ELISPOT in splenocyte populations stimulated with peptide pools (15AA length with 11mer overlaps) spanning the spike protein 90 days after vaccination. (**f**) Intracellular cytokine staining and FACS analysis showing frequency of IFN-γ^+^ CD8^+^ T cells specific for RBD region of the spike protein. (**g**) Pie charts comparing the mean proportions of single, double, and triple cytokine producing cells (left panels) and frequency of multifunctional IFN-γ-producing T cell populations (right panels). (**h**) Absolute numbers of RBD-specific triple cytokine (IFN-γ^+^ TNF ^+^ IL-2^+^) cells within the cytokine positive CD8 T cell compartment. Symbols represent individual mice and bars show the mean ± SEM. Data was log transformed and statistical significance was determined using Brown-Forsythe and Welch ANOVA with Dunnett T3 multiple comparison test. *p<0.05; ***p<0.01; ***p<0.001; ****p<0.0001

Next, the impact of pre-existing immunity on spike-specific memory CD8 T cell responses at three months post SCV-S vaccination was evaluated in the spleen using peptide pools specific to S1-NTD, RBD, and S2 regions of the spike-protein. As anticipated, irrespective of the pre-existing immunity status, SCV-S vaccinated mice had significantly higher S1-NTD-, RBD-, and S2-specific IFN-γ^+^ SFU compared to control vaccinated mice (Figure 5e). However, approximately 10-15-fold lower S1-NTD- and RBD-specific T cell responses were noted in mice with pre-existing VACV memory responses when compared to VACV naïve mice, suggesting the magnitude of the spike-specific T cell responses was impacted on by pre-existing vector immunity. Assessment of the IFN-γ^+^ CD8^+^ T cell population confirmed a 20-fold increase in IFN-γ expression in response to SCV-S prime-boost vaccination in VACV naïve mice compared to VACV experienced cohorts (Figure 5f).

Further analysis of the dominant RBD-specific CD8^+^ T cell responses revealed a higher proportion of RBD-specific multifunctional T cells in VACV naïve mice compared to VACV experienced mice, with a specific increase in IFN-γ^+^ cells co-expressing TNF (Figure 5g). Despite a lower proportion of multifunctional T cells within the cytokine producing CD8^+^ T cells, significant numbers of RBD-specific triple-cytokine positive cells in VACV experienced mice could still be detected compared to control vaccinated mice (Figure 5h). Together this indicated that pre-existing vector immunity had no significant effect on the humoral immune response, and whilst significantly reduced the magnitude of the spike-specific memory CD8^+^ T cell response, vaccination with SCV-S could still however induce significant spike-specific cellular responses.

### Aging mice demonstrate long-lived and robust spike-specific cellular and humoral immune responses following SCV-S vaccination

Long-lived immunological memory forms the basis of robust prevention of disease following vaccination. Previous studies have demonstrated that impaired immune responses in older individuals can be associated with decreased immunological memory post-vaccination^39–41^. Therefore, the ability of SCV-S to establish and retain a long-lived spike-specific immune response was investigated. Young and aging C57BL/6J mice were vaccinated in a 4-week prime-boost regimen, and spike-specific immune responses were analysed nine months post-vaccination. Significant S1 binding IgG titers were observed in both young and aging mice compared to control vaccinated mice, with no statistical difference observed between the groups (Figure 6a, left panel). Surprisingly, S2 binding titers were detected only in 4 out of 5 aging mice, and while the mean S2 IgG binding titers in the aging mice were higher than the control vaccinated mice, it was not statistically different. The young mice on the other had significant S2 IgG binding titers compared to the control vaccinated mice (Figure 6a; right panel). Retrospective analysis of the S2 binding titers in the young and aging mice during the vaccination time course revealed that the levels were comparable at day 21 post prime, however no further increase in the S2 binding titers in aging mice followed the booster dose. In comparison, an approximate 10-fold increase in the S2 binding titers were detected in young mice after boost vaccination (Supplementary Figure 9). Despite these differences in the S2 binding titers, the neutralisation activity was comparable between the young and aging mice at 9 months (Figure 6b). Persistent antigen-specific antibody levels in the serum are maintained by long-lived antibody secreting cells (ASC) that reside in the bone-marrow^42–44^. Therefore, the capacity of SCV-S prime-boost vaccination to induce a S1-specific ASC population by B cell ELISPOT was investigated. Both the young and aging mice demonstrated significant numbers of S1-specific ASCs compared to the control vaccinated mice, with the populations comparable between the two groups (Figure 6c).

**Figure 6:**
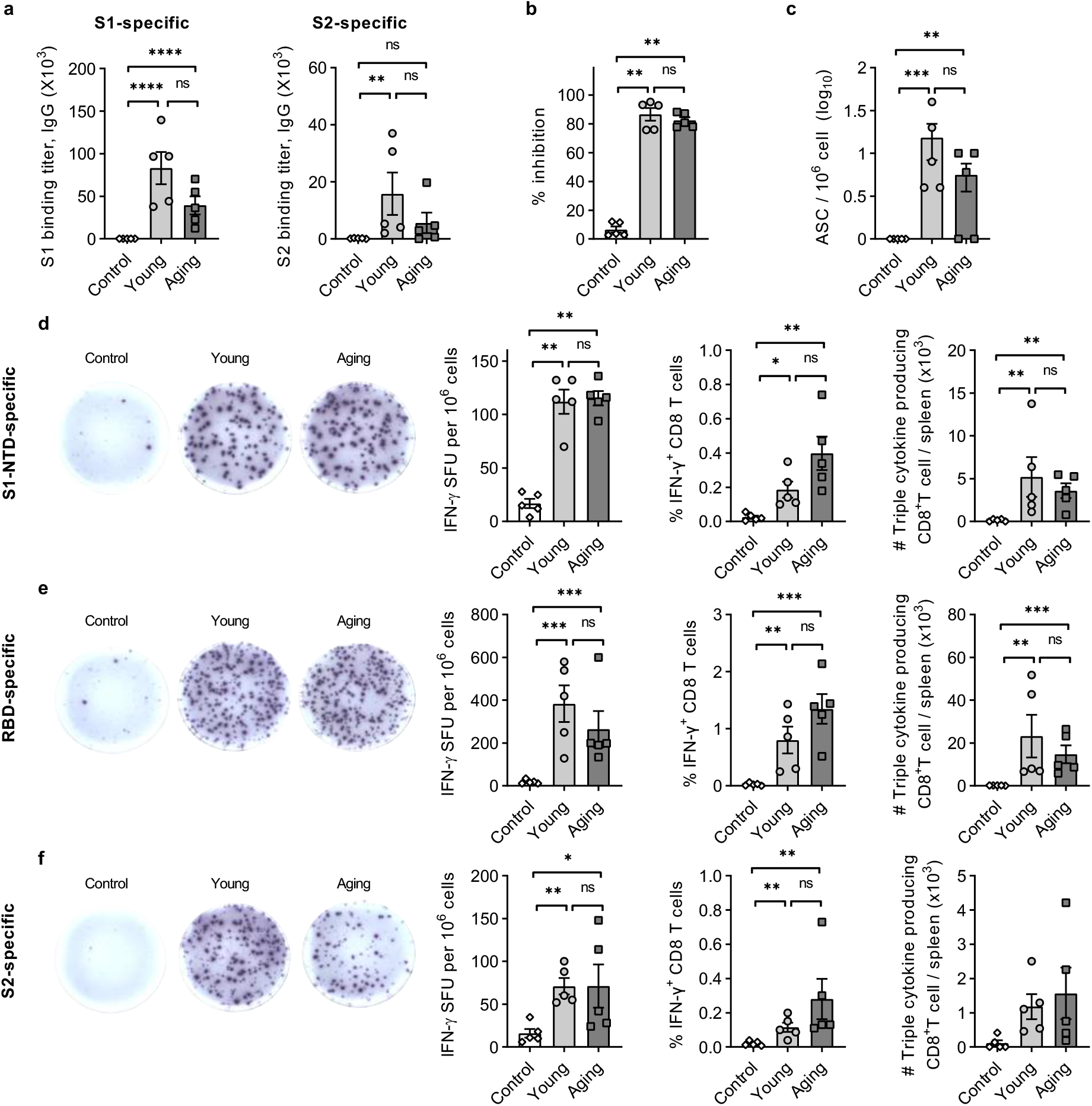
Longevity of spike-specific B and T cell responses following SCV-S vaccination. Cohorts of young (6-8 weeks old) and aging (9-10 months old) female C57BL/6J mice (n=5) were vaccinated in a prime-boost regimen (day 0 and 28) with 10^7^ PFU of SCV-S or control vector. (**a**) S1-specific (left panel) and S2-specific (right panel) endpoint IgG ELISA titers and (**b**) neutralization titers 9 months after vaccination. (**c**) Frequency of long-lived S1-specific IgG antibody secreting cells (ASC) assessed by B cell ELISPOT. **(d,e,f)** Peptide pools (15AA length with 11mer overlaps) spanning the spike protein were used to stimulate splenocytes 9 months after vaccination and IFN-γ SFU specific for S1-NTD, RBD, and S2 subunit regions of the spike protein were quantitated by ELISPOT (left panels), with frequency of IFN-γ^+^ CD8^+^ T cells (middle panels) and absolute numbers of triple cytokine producing cells (right panels) enumerated by intracellular cytokine staining and FACS analysis. Symbols represent individual mice and bars show the mean ± SEM. Data was log transformed and statistical significance was determined using Brown-Forsythe and Welch ANOVA with Dunnett T3 multiple comparison test. **p<0.05; ***p<0.01; ***p<0.001

Consistent with the antibody responses, spike-specific T cell responses were maintained in both young and aging mice at nine months post SCV-S vaccination. Significant numbers of S1-NTD (Figure 6d), RBD (Figure 6e) and S2 (Figure 6f) specific IFN-γ^+^ SFU and IFN-γ^+^ CD8^+^ T cells were detected in both the young and aging mice compared to the control vaccinated mice, with no statistical difference noted between the two groups. The proportions of multifunctional T cells within the cytokine producing CD8^+^ T cell population across all three regions of the spike protein were also comparable (Supplementary Figure 10). Triple cytokine producing CD8^+^ T cells which have enhanced cytokine and memory potential were also detected in significant numbers in all groups except S2 targeting triple cytokine producing CD8^+^ T cells which suggested that S1-specific polyfunctional effector cellular immune responses dominate the memory compartment following SCV-S vaccination. In summary, these results indicate that SCV-S vaccination induces a long-lived humoral and cellular immunity, with similar magnitude and quality of immune responses in both young and aging mice, demonstrating broad applicability of SCV-S.

## Discussion

Vaccine development programs that enhance existing vaccine technologies and advance development of novel vaccine platforms are critical in the current race to curb SARS-CoV-2 evolution and control the COVID-19 pandemic. Complementary and synergistic vaccination strategies and cohort-specific vaccination approaches are rapidly gaining acceptance, due to a lack of convenient and effective antiviral therapies, a broad range of vulnerable populations, and the intrinsic manufacturing challenges associated with global vaccination campaigns. In the current stage of the pandemic, a potential vaccine candidate should satisfy one or more of the following criteria: induce a broad and robust humoral and cellular immune response against the dominant circulating variants either as a stand-alone or effective booster to approved vaccines, enhance immunity in vulnerable populations, establish long-lived immune responses that can maintain herd immunity and prevent circulation of SARS-CoV-2 and variants, provide cross-protection against other coronaviruses, and have a stream-lined and facile manufacturing process.

The SCV vaccine platform technology builds on the favourable characteristics of the parental VACV vector, namely long-lasting cellular and humoral immune responses, significant antigen payload load capacity, cold-chain-independent vaccine distribution capability, and incorporates additional advantages primarily safety while maintaining immunogenicity. SCV-based vaccines have proven stability at 4°C for 6 months in simple salt buffered liquid formulations, and at least 1 year in dried formulations (unpublished data) which may address the logistical difficulties associated with vaccine deployment in hard to reach and vulnerable communities. Furthermore, the SCV platform addresses the manufacturing challenges associated with the use of primary cells in empirically attenuated VACV-based vaccine production; industry-standard CHO cells were uniquely genetically engineered to allow high yield production of SCV vaccines. This primary study describes the immunogenicity of a first-generation SCV-mediated COVID-19 vaccine. SCV-S incorporates a single antigen of the SARS-CoV-2, the spike glycoprotein, that mediates viral attachment and entry into host cells. The data presented here shows that SCV-S infected cells produce spike immunogen that is transported from the endoplasmic reticulum through the Golgi apparatus to the cell surface to stimulate antigen-specific immune responses. Importantly, the spike insert was maintained in the viral genome without any loss or changes in transgene sequence or protein expression, thus confirming the genetic stability of the vaccine through several passages, which is an inherent characteristic to support large-scale vaccines manufacturing protocols.

Immunogenicity studies in mice confirmed that the vaccine induces spike-specific functional T cell responses in 1 week, and robust levels of circulating spike-specific antibodies in both inbred and outbred strains of mice, with significant neutralizing activity detected within two weeks of vaccination. These functional immune responses satisfy the rapid onset of protection criteria set forth in the COVID-19 vaccine target product profile by WHO, a promising attribute for a stand-alone vaccine in establishing ring immunity and preventing spread of infection. Whilst a direct comparison of the neutralising activity with other front-runner vaccines would be useful, differing protocols and vaccine assessment regimens used across diverse vaccine platforms deployed and in development make it virtually impossible to do so. Instead, neutralizing activity against SARS-CoV-2 and the VOCs (Alpha and Beta) was confirmed by two different methods, (1) using an FDA approved validation methodology^45^ that detects antibodies that block the interaction between RBD (and variants thereof) and the ACE2 cell surface receptor, and (2) a lentiviral based pseudovirus neutralization assay. Importantly, after two doses the circulating neutralizing antibody responses were maintained long-term with minimal contraction, up-to nine months post-vaccination, in both young and aging mice. The sustained circulating antibody levels are encouraging particularly in light of recent studies suggesting a decay in neutralisation activity in vaccinated^46^ and convalescent patients^47^, and support protective efficacy studies in infectious disease challenge models and further development to establish safety and efficacy of future SCV-based COVID-19 vaccines in human clinical trials.

To date, the correlates of protection against SARS-CoV-2 in humans has not been clearly defined. Whilst it is well acknowledged that neutralizing antibodies play a critical role, emerging clinical data suggest the presence of SARS-CoV-2 specific T-cell responses may be an important differentiating factor between levels of resultant COVID-19 severity^48^. Preclinical efficacy studies in mouse models have demonstrated that in the absence of antibodies, T cell responses can protect against SARS-CoV-2 infection^49^. In addition, recent studies show that the SARS-CoV-2 VOCs can partially escape broadly neutralizing antibody responses but maintain effector and memory T cell reactivity^50^, further highlighting the complimentary role of T cells in modulating the disease severity for emerging variants. Following vaccination with SCV-S, robust spike-specific polyfunctional CD8 T cells with cytotoxic activity was observed within the first week, which further differentiate to form a memory T cell pool that can be detected upon secondary stimulation nine months post-vaccination, in both young and aging mice. Notably, spike-specific humoral and cellular immune responses could be detected even in the presence of pre-existing vaccine vector immunity.

Analysis of the antibody repertoire in SARS-CoV-2 recovered individuals revealed the presence of neutralizing antibodies against the S1-NTD, and S2 region of the spike protein in addition to anti-RBD antibodies^51, 52^. Whilst the exact role and relative contributions of the non-RBD neutralizing antibodies in SARS-CoV-2 infection remains to be fully addressed, it is intuitive to deduce that broad spike-specific antibody responses spanning mutational hotspots (RBD and S1-NTD) and conserved regions (S2) of the spike region, *via* a combination of anti-S1 and anti-S2 neutralizing antibodies, will offer improved protective efficacy potential against emerging variants. T cell epitope analysis in naïve individuals revealed reactivity to SARS-CoV-2, with significant proportion of the epitopes mapped to the non-RBD (44%) region of the spike protein^53^, suggesting that T cell responses against the conserved S2 region can offer cross-protection against other betacoronaviruses. Our results demonstrate that SCV-S induces a broad-spike specific antibody and CD8 T cell response spanning both the S1 and S2 subunits, enhancing potential activity against other betacoronaviruses and emerging novel SARS-CoV-2 variants.

In summary, the data presented in this study demonstrate that the SCV-S vaccine induces a rapid, broad, robust, and durable spike-specific cellular and humoral response supporting the progression of SCV-based COVID-19 vaccine candidates towards authentic efficacy testing in recognised preclinical SARS-CoV-2 infectious disease challenge trials, and clinical development programs. Furthermore, poxvirus-based vectors have a long history as efficient boosters to a wide range of platform technologies such as DNA^54^, protein^55^ and viral vectored platforms^56^. As such, the evidence of serological and cellular priming by the SCV-S vaccine in this study, along with the safety profile of the replication-defective SCV platform, suggest that the vaccine will also be an effective booster to the current vaccines, with potential to enhance spike-specific immune responses. The encouraging immunogenicity profile with a single SARS-CoV-2 antigen also provides impetus to include additional SARS-CoV-2 antigens in second-generation SCV-COVID-19 vaccines, primarily to broaden and strengthen vaccine immunogenicity profiles for enhanced vaccine protective efficacy against current VOCs and new emerging variants as they inevitably become established within the affected global population.

## Materials and methods

### Cell lines

The CHO-based manufacturing cell line (MCL) expressing D13 and CP77^25^ was maintained in CD-CHO media supplemented with 8 mM L-glutamine, penicillin-streptomycin (1 U/ml and 0.1 mg/ml, respectively) solution, hygromycin B (500 μg/ml), and puromycin (10 μg/mL). 143B cells (ATCC CRL-8303) were maintained in RPMI-1640 culture media supplemented with 10% (v/v) fetal bovine serum (FBS), 2 mM L-glutamine, and penicillin-streptomycin. Virus titration was performed in 143B cells expressing D13 (ST01-33) maintained in 143B culture media supplemented with hygromycin B (500 μg/ml). MC57G cells (ATCC CRL-2295) for chromium release assays were maintained in Eagle’s Minimum Essential medium supplemented with 10% FBS, 2 mM L-glutamine, and penicillin-streptomycin. ACE2-HEK293 recombinant cell line (Cat # 79951; BPS Bioscience) for use in pseudovirus neutralization assays were maintained in Dulbecco’s Minimum Essential medium supplemented with 10% FBS, 2 mM L-glutamine, and penicillin-streptomycin.

### Animal experiments

Specific-pathogen-free inbred C57BL/6J and outbred Swiss mouse heritage Arc:Arc(S) mice were bred in house or purchased from the Animal Resources Centre (Canning Vale, WA, Australia). All experiments were conducted under protocols approved by the University of South Australia Animal Ethics Committee in accordance with the Australian code for the care and use of animals for scientific purposes, 8^th^ edition (2013). Vaccinations were performed under isoflurane anaesthesia by intramuscular administrations into both quadricep muscles of the hind legs. Serum samples obtained by routine blood collection were stored at -80°C until further analysis.

### Construction of SCV-S vaccine

SCV-S vaccine was constructed by replacing the A41L ORF with the SARS-CoV-2 spike glycoprotein sequence (21563-25384 bp gene from Wuhan-1 isolate; Genbank accession # MN908947) by homologous recombination. The SARS-CoV-2 spike ORF was synthesised (GeneArt; Thermo Fisher Scientific) with a poxvirus early transcriptional stop sequence (T5NT) under the control of a synthetic vaccinia virus early/late promoter and cloned into the *Pac*I site of a transfer plasmid containing a comet GFP fused with the Zeocin resistance gene expression cassette flanked by a 150bp repeat sequence for intramolecular recombination, and F1 and F2 arms homologous to upstream and downstream sequences of the *A41L* ORF using in-fusion cloning (Takara Bio). The resultant spike transfer cassette was linearized by *Not*I and transfected into the MCL for homologous recombination with SCV, a recombinant version of VACV in which *D13L*, *B7R/B8R*, *C3L*, and *A39R* were previously deleted, at an MOI of 0.01. Five rounds of single cell amplification in the MCL was performed to obtain recombinant clones which were subsequently amplified in the absence of Zeocin to promote intramolecular recombination and deletion of the fluorescent reporter cassette. Non-fluorescent cells were amplified by bulk sorting and the research grade master virus seed (MVS) stocks was generated for vaccine characterisation.

### Vaccine stocks and titration

Vaccine stocks were prepared by infecting MCL cells (MOI of 0.01) maintained at 33°C, 5% CO_2_ in 1L baffled Erlenmeyer flasks shaking at 110 rpm for 48hrs in CD-CHO media supplemented with 8mM L-glutamine. Cells were recovered by centrifugation at 2000 *x* g for 10 minutes at 4°C. To release the virus from the cells, the harvested cell pellet was incubated with lysis buffer (1% Tween-80 in 10mM Tris-HCl, pH 8) for 30 mins at room temperature (RT) with constant mixing. The clarified supernatant (1000 *x* g for 10mins at 4°C) was layered onto a 36% sucrose cushion and centrifuged at 3200 *x* g for 18hr at 10°C to pellet the virus. Vaccine stocks were resuspended in 10mM Tris-HCl and stored at -80 °C until further use. Vaccine titers were determined by plating serial dilutions of the vaccine stocks on 80% confluent ST01-33 cells in 24 well plates, incubation at 37 °C, 5% CO_2_ for 48 hrs and plaque staining with crystal violet. Titers were expressed as PFU/mL.

### *In vitro* characterisation of SCV-S vaccine

Construction and purity of the SCV-S MVS stock was confirmed by polymerase chain reaction (PCR) using primers spanning the inserted and deleted regions of the viral genome. Genomic DNA from infected MCL cells was extracted using the NucleoSpin Tissue kit (Macherey-Nagel). PCR reactions were performed using KAPA HiFi polymerase (KAPA biosystems) using the following primer pairs: D13L locus, forward primer 5’-GTGAGTACCCTGGATACGAAATAAA-3’ and reverse primer 5’-AACCATCTACAGTATACGTTTATATTAAAA-3’; A41L locus, forward primer 5’-TCACATCGTTTACGCAATAGTCAGACT-3’ and reverse primer 5’-CTAGACGAACCCCTCAGACAAACAAC-3’ and spike insertion primers, forward primer 5’-TATTCCAGTTAAAGCACGGTTTAATTGTGTAC-3’ and reverse primer 5’-GTTACCAACCATACAGAGTAGTAGTACTTTCTTTG-3’. PCR products were run on 1% agarose gel and visualised using GelRed (Biotium) and transgene sequence confirmed by Sanger sequencing at the Australian Genome Research Foundation (AGRF).

To assess the genetic stability of SCV-S, serial passages of the MVS were performed in triplicate. On the day of infection, cells were resuspended in 20mL of fresh growth media at 10^6^ cells/ml and infected at MOI 0.01 and incubated shaking at 33°C. On day 3 post-infection, cells were harvested by centrifugation at 300 *x* g for 5 minutes and resuspended in 1 mL of 10 mM Tris, pH 8. A homogenate was generated using a bead homogeniser (MP biomedicals) and the viral titer of the clones was determined by plaque assay to enable the next infection to be performed in the same manner. Genomic DNA was extracted for Illumina NGS from 100 mL cultures of MCL infected at MOI 0.1. Cells were harvested and lysed using 1% Tween-80 in 10mM Tris-HCl, pH 8. Nuclei and other cell debris was cleared by centrifugation and the remaining viral lysate was incubated overnight with benzonase (25-50 U/ml) to further degrade the host cell DNA. Virus was purified by centrifugation at 12,000 xg for 10 mins at 4 °C and genomic DNA extracted (NucleoSpin Tissue kit, Macherey-Nagel). Next generation sequencing (NGS) was performed using the Illumina platform (South Australian Genomics Centre, Adelaide). Paired-end reads (150bp) were mapped to the reference sequence and variants examined using CLG Genomics Workbench 20 software.

### Western blot analysis of spike protein

Cell lysates were collected from 143B cells after 24hr infection with SCV-S or control vector with proteins separated by electrophoresis (10% SDS-PAGE), transferred to nitrocellulose membrane, blocked with 5% skim milk in PBS-T and incubated with rabbit SARS-CoV-2 RBD polyclonal antibody (Cat # 40592-T62, Sino Biological; 1:1,000 dilution) overnight at 4°C. Following washing, the membrane was incubated with anti-rabbit IgG antibody conjugated to horseradish peroxidase (Cat # A9044, Sigma Aldrich; 1:10,000 dilution) for 1 hr at RT and proteins visualized using Clarity ECL western blotting substrate (BioRad) and imaged using ChemiDoc XRS (BioRad).

To examine spike expression following serial passage, the MCL was infected (MOI of 0.01) for 48 hrs, harvested, titrated, and used for infection of next passage. A total of 10 rounds of low MOI passaging was performed, with passage 10 (P10) stock used to infect 143B cells for immunoblot analysis.

### Spike protein expression by flow cytometry

To confirm processing of the spike protein and surface translocation, 143B cells were infected with SCV-S or SCV control (empty) vector with an MOI of 1 or left uninfected. Cells were harvested at 24 hrs, stained with live-dead marker (LIVE/DEAD fixable e660 stain, Thermo Fisher Scientific; 1/3000 dilution) and anti-SARS-CoV-2 RBD antibody (Cat # 40592-T62, Sino Biological; 1:100 dilution), followed by staining with PE-conjugated donkey anti-rabbit IgG (Cat # 406421, Biolegend), data was acquired using a FACSARIA Fusion™ (BD Biosciences), and analysis performed using FlowJo V10 software.

### Plaque immuno staining

Monolayers of ST01-33 in 24 well plates infected with virus (control, SCV-S MVS or SCV-S P10 vaccine stocks) were fixed after 48 hrs with 1:1 solution of methanol/ acetone and air-dried prior to immunostaining with rabbit SARS-CoV-2 RBD polyclonal antibody (Cat # 40592-T62; Sino Biological; 1:1000 dilution) and HRP-conjugated anti-Rabbit IgG antibody (1:2000; Life Technologies) and developed using TMB substrate for membranes (Sigma-Aldrich).

### Infectivity assays

To compare the replication capacity of MVS and P10 vaccine stocks, 30 mL cultures of MCL cells were infected at an MOI of 0.01, harvested at 0, 24,48,72 and 96 hrs post-infection and titrated on ST01-33 monolayers to determine the fold increase in titer.

### IFN-γ ELISPOT assay

Antigen-specific IFN-γ producing T cell responses post-vaccination were examined in the spleen. ELISPOT plates (MSIPS4510; Millipore) were activated with 15% ethanol, washed, coated with anti-mouse IFN-γ capture antibody (clone AN18; Thermo Fisher Scientific; 8 μg/mL in PBS) and incubated overnight at 4°C. The next day, single cell suspensions of splenocytes were prepared, serial diluted into blocked ELISPOT plates, and stimulated with a final concentration of 2 μg/mL of each peptide for 18 hrs at 37°C in a humidified 5% CO2 incubator. Plates were then washed and incubated with biotinylated anti-mouse IFN-γ antibody (clone R4-6A2; Mabtech; 1 μg/mL) followed by streptavidin-alkaline phosphatase (Mabtech; 1:1,000 dilution). Spots were visualised using BCIP/NBT plus substrate (Mabtech) and counted using an automated ELISPOT plate reader (AID vSpot Spectrum).

### Cytotoxic T-lymphocyte (CTL) assay

Spike-specific CTL activity was assayed using standard chromium release assay. Briefly, MC57G target cells were pulsed with 2 μg/mL of each peptide for 2 hrs and then labelled with ^51^Cr for 1hr. Single cell suspensions of effector splenocytes were then incubated with MC57G target cells for 6 hours starting at an effector: target cell ratio of 100:1. For each peptide pulse condition, maximum and spontaneous release conditions were set up in sextuplicate. Plates were centrifuged and 30 μL of the supernatant transferred to a 96-well Luma plate (Perkin-Elmer), allowed to dry overnight, and counts analysed on a top count Microbeta plate reader. Percent specific lysis was determined using the equation: [(Sample ^51^Cr release − Spontaneous ^51^Cr release)/(Maximum ^51^Cr release − Spontaneous ^51^Cr release)] ×100.

### T cell analysis by flow cytometry

Intracellular cytokine staining was performed by stimulating splenocytes with a final concentration of 2 μg/mL per peptide for 2 hrs at 37°C. Brefeldin A (final concentration 10 µg/mL) was subsequently added for an additional 4 hrs. The cells were washed in FACS buffer (PBS+ 2% heat-inactivated FCS) and surface stained for 30 mins at 4°C with 50 μL/well anti-mouse CD3 APC-CY7 (clone 17A2, 1:400, BD Biosciences), anti-mouse CD4 BV510 (clone RM4-5, 1:400, BD Biosciences) and anti-mouse CD8a PE (clone 53-6.7, 1:400, BD Biosciences). Cells were washed with PBS and incubated with LIVE/DEAD fixable e660 stain, for 20 min at 4°C. Cells were washed with PBS and then fixed and permeabilised using Cytofix/cytoperm kit (BD Biosciences) as per manufacturer’s instructions. Intracellular cytokines were stained with anti-mouse IFN-γ FITC (clone XMG1.1, 1:200, BD Biosciences), anti-mouse TNF-α BV421 (clone MP6-XT22, 1:200, BD Biosciences), anti-mouse IL-2 PE-CF594 (clone JES-5H4, 1:200, BD Biosciences), and anti-human/mouse granzyme B PE-Cy7 (clone Q1A6A02, 1:200, BioLegend). All antibodies were diluted in Brilliant stain buffer (BD Biosciences) and added to cells resuspended in permeabilization buffer and incubated overnight at 4°C. Cells were washed, resuspended in 150 µL FACS buffer, and results acquired on FACSARIA Fusion™ with analysis by FlowJo V10 software.

Memory T cell populations were identified by staining for surface markers with the following anti-mouse antibodies (BD Biosciences): CD3 APC-CY7 (clone 17A2, 1:400); CD4 BV510 (clone RM4-5, 1:400); CD8a PE (clone 53-6.7, 1:400); CD44 FITC (clone IM7,1:300, BD Biosciences); KLRG1 PE-CF594 (clone 2F1, 1:400); CD62L BV650 (clone MEL-14,1:400); CD127 BV421 (clone A7R34, 1:400).

### Binding antibody titers

S1-, S2- and vector-specific binding antibody titers were quantitated by enzyme linked immunosorbent assay (ELISA). High-binding 96-well plates (Nunc) were coated with the appropriate antigen (S1 protein 1.2 μg/ml; S2 0.8 μg/mL; SCV 10^5^ PFU/well) in carbonate buffer (pH 9.8) and incubated at 4°C overnight. Plates were washed with PBS-T, blocked with 5% skim milk powder, and then incubated with 3 fold dilutions of relevant serum samples (starting dilution 1:100) in 1% skim milk powder in PBS-T for 2hrs at RT. Binding serum antibodies were detected using HRP-conjugated anti-mouse IgG (Cat #A9044, Sigma Aldrich, 1:10,000), IgG1 (Cat #1070-05, Southern Biotech, 1:1,000), IgG2b (Cat #1090-05, Southern Biotech, 1:1,000), IgG2c (Cat #1079-05, Southern Biotech, 1:1,000), IgG3 (Cat #11100-05, Southern Biotech, 1:1,000), with colorimetric signals developed with 100 μL/well of 3,3ʹ,5,5ʹ-Tetramethylbenzidine (TMB) Liquid Substrate (Cat # T0440, Sigma-Aldrich) and stopped using 50 μL/well of 3M HCl. Absorbance readings were acquired at 450 nm wavelength. End-point titers were defined as the highest reciprocal of serum dilution to yield an absorbance equal to the negative serum samples plus three times the standard deviation.

### SARS-CoV-2 neutralization antibody detection using cPass™ kits

Levels of neutralizing antibodies in serum samples were determined by cPASS^TM^ SARS-CoV-2 neutralization antibody detection kit (GenScript, USA) as per the manufacturer’s instructions. Briefly, serum samples and controls were diluted 1:10, incubated with HRP-conjugated RBD for 30mins, and transferred to a capture plate coated with ACE2 protein and further incubation for 15mins. Plates were washed, with activity detected using TMB substrate. Plates were read at 450 nm and percent signal inhibition determined using the formula: Percent Signal Inhibition = (1 – OD value of sample / OD value of negative control) x 100%

### Lentivirus-based pseudovirus neutralization assay

Neutralization antibody activity against SARS-CoV-2 variants was examined using spike-pesudotyped lentiviruses in ACE2-HEK293 recombinant cell line and quantitated as reductions in luciferase reporter expression as per manufacturer’s instructions (BPS Bioscience). Original SARS-CoV-2 spike pseudotyped lentivirus (Cat # 79442), spike B.1.1.7 (Alpha) variant pseudotyped lentivirus (Cat # 78112) and spike B.1.351 (Beta) variant pseudotyped lentivirus (Cat # 78142) were used. ACE2-HEK293 cells were plated at a density of 10,000 cells/well in a clear bottomed white opaque 96-well plate and incubated at 37°C. The next day, three-fold serial dilutions of the serum samples were incubated with the pseudoviruses for 30mins and plated onto ACE2-HEK293 cells. After 24hrs, the media was changed, and the following day cells were lysed using ONE-Step™ Luciferase kit and luciferase activity was measured. Data was normalised as percent neutralization using the readings from uninfected cell controls to set the baseline for 100% neutralization and the pseudovirus alone controls to set the baseline for 0% neutralization. IC_80_ titers were determined using a log (inhibitor) vs. normalized-response (variable slope) nonlinear regression model in Prism v9 (GraphPad).

### Antibody secreting cell ELISPOT

S1-specific IgG producing antibody secreting cells in bone-marrow was examined by ELISPOT assay. ELISPOT plates (MSIPS4510; Millipore) were activated with 15% ethanol, washed, coated with S1 protein (10 μg/mL in PBS) and incubated overnight at 4°C. The next day, harvested bone-marrow cells were resuspended as 5x 10^6^ cells/mL and 100µL added to washed and blocked ELISPOT plates for 24 hr culture at 37°C/5% CO_2_. Plates were subsequently washed, incubated with biotinylated anti-IgG detection antibody (Mabtech, 1 μg/mL), followed by streptavidin-alkaline phosphatase (Mabtech, 1:1,000 dilution). Spots were visualised using BCIP/NBT plus substrate and counted via ELISPOT plate reader.

### Statistics

GraphPad Prism version 9.0 (GraphPad software) was used for data analysis and statistics. Data was log-transformed and where appropriate Brown-Forsythe and Welch ANOVA with Dunnett T3 multiple comparison test for unpaired samples or repeated measures ANOVA with Tukey multiple comparison test for paired samples was used to determine statistical significance between three or more group. For statistical analysis between two groups, *t*-tests with Welch’s correction was used. Correlation analysis was performed using Spearman rank test.

## Data availability

All datasets generated and analysed in the current research are available from the corresponding authors upon reasonable request.

## Acknowledgements

The authors acknowledge the contributions of Paul M. Howley, past CEO/CSO of Sementis Ltd., and Robyn Kievit for technical support to this work, as well as The University of Queensland Protein Expression Facility and Genescript for providing reagents and additional support.

## Author Contributions

P.E., T.C., N.P., L.L., R.G.G, P.W., K.D. and J.H. designed experiments; P.E., T.C., N.P. and L.L. constructed the vaccine and along with GH, JZ, and AT acquired the data. P.E. and K.D. analysed the results, prepared the figures, and wrote the manuscript. All authors discussed the results and edited the manuscript. J.H., K.D., and L.H. supervised the study.

