## Supplementary information for "The vaccinia-based Sementis Copenhagen Vector COVID-19 vaccine induces broad and durable cellular and humoral immune responses"

**Supplementary Table 1:** Peptide pools spanning the spike glycoprotein used in T cell analysis

| Peptide pool nomenclature | Amino acid no. | Region | # of peptide pooled |
|---|---|---|---|
| S1-NTD | 1-318 | 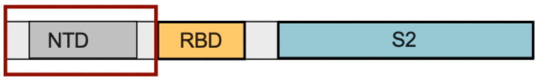 | 77 |
| RBD | 319-685 | 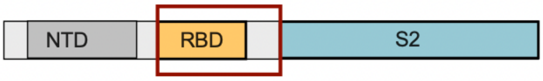 | 92 |
| S2 | 686-1273 | 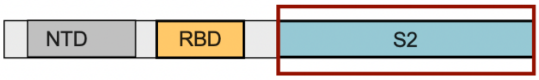 | 146 |

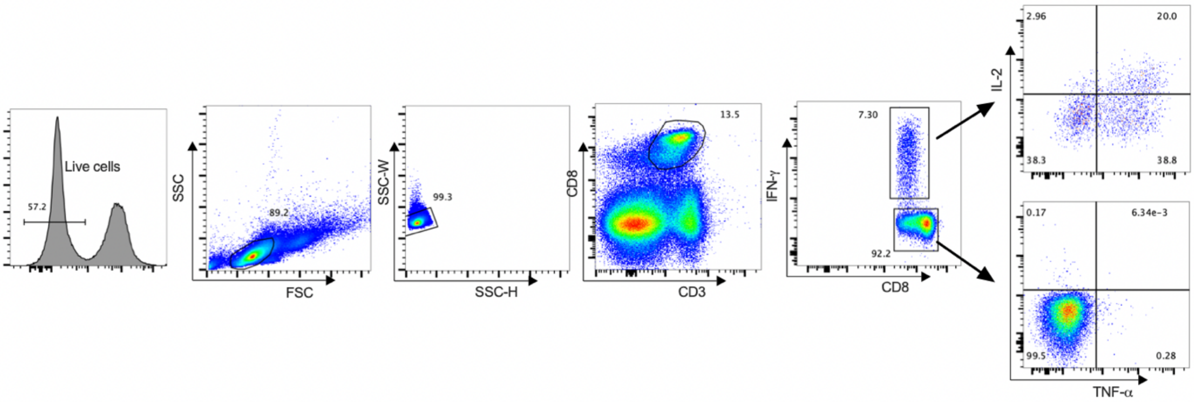


**Supplementary 1: Gating strategy for spike-specific multifunctional T cells.**
C57BL/6J female mice (n=5) were vaccinated IM with $10^7$ PFU of SCV-S. At day 7 post-vaccination, splenocytes were pulsed with peptide pools (15AA length with 11mer overlaps) spanning the S1 subunit of spike protein and viable single $CD3^+$ $CD8^+$ T cells were enumerated for intracellular cytokine staining using the gating strategy shown. The numbers within the gates indicate the percentage of parent population.

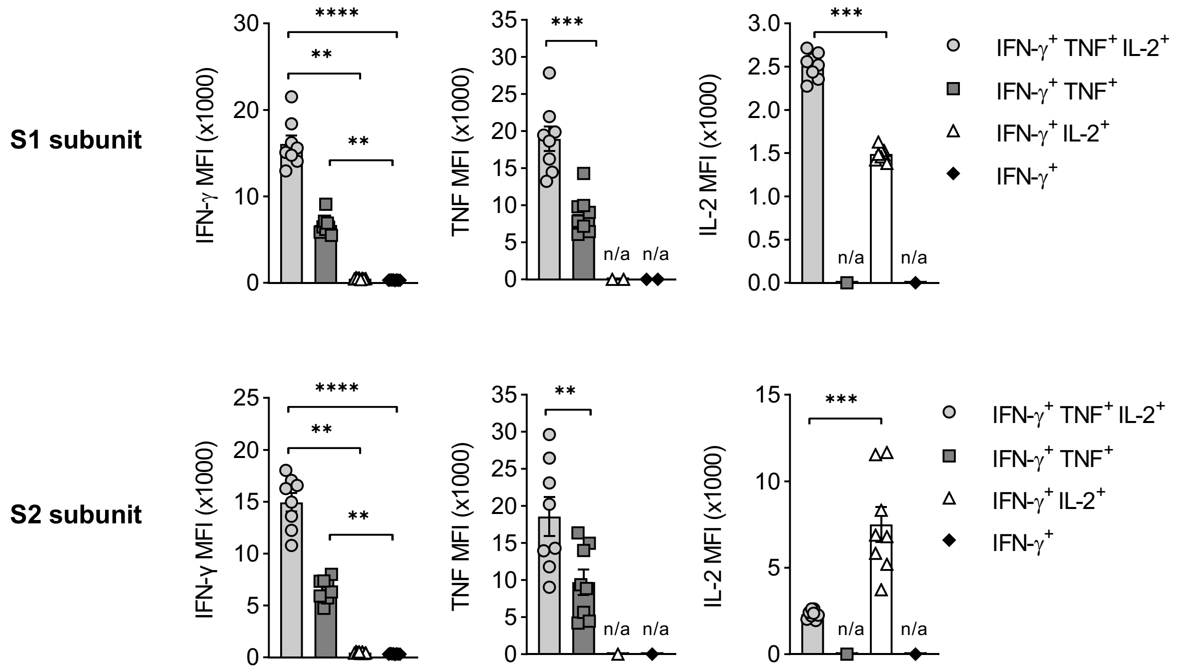


**Supplementary 2: Magnitude of cytokine production in spike-specific $CD8^+$ T cell populations.**

C57BL/6J female mice (n=5) were vaccinated IM with $10^7$ PFU of SCV-S. At day 7 post-vaccination, splenocytes were pulsed with peptide pools (15AA length with 11mer overlaps) spanning the S1 and S2 subunits of the spike protein and intracellular cytokine production assessed by flow cytometric methods. The magnitude of IFN-γ, TNF, and IL-2 was measured by median fluorescence intensity (MFI) in single, double, and triple cytokine producing $CD8^+$ T cells specific for the S1 and S2 subunits. Symbols represent individual mice and bars show the mean ± SEM. Statistical significance was determined using Kruskal-Wallis ANOVA test or Mann-Whitney t-test, $^{**}p<0.01$; $^{***}p<0.001$; $^{****}p<0.0001$, n/a=not applicable

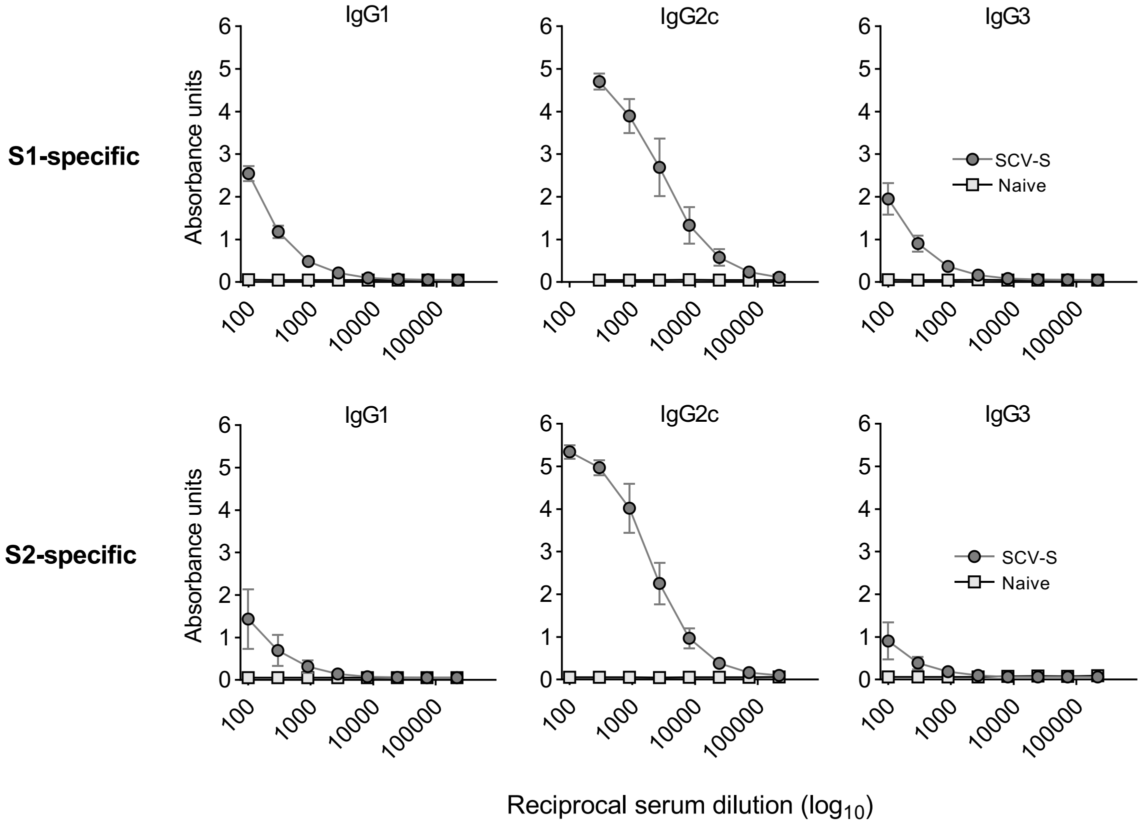


**Supplementary 3: Subclass profiling of spike-specific antibody responses.**
C57BL/6J female mice (n=3) were vaccinated with $10^7$ PFU of SCV-S or left untreated. At day 21 post-vaccination, subclass profiling of S1- and S2-subunit specific antibody responses was assessed by ELISA to establish Th1 bias.

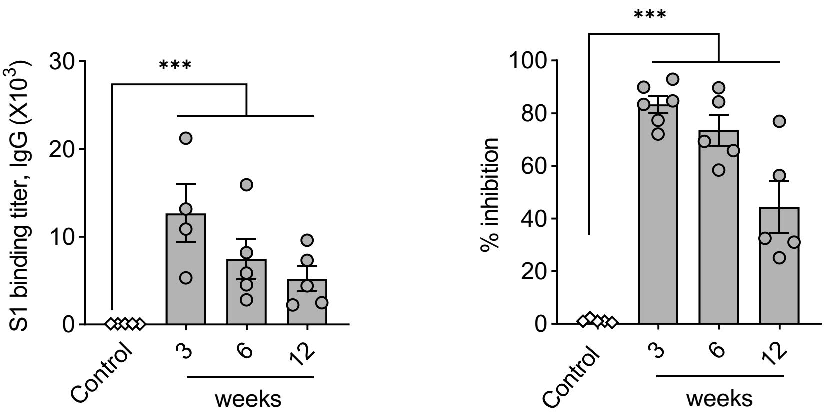


**Supplementary 4: Maintenance of spike-specific antibody responses in outbred mice following single SCV-S vaccination.**

Inbred C57BL/6J female mice (n=5) were vaccinated with $10^7$ PFU of SCV-S and serum samples collected at times indicated. S1-specific antibody binding was assessed by ELISA (left panel), with endpoint IgG titers reported (middle panel). Neutralizing titers were determined by cPASS SARS-CoV-2 neutralization antibody detection kit (right panel). Symbols represent individual mice and bars show the mean ± SEM. Repeated measures one-way ANOVA with Geisser-Greenhouse correction was used to compare the samples with Fisher's LSD test used to report multiple comparisons. *p<0.05; **p<0.01; ***p<0.001

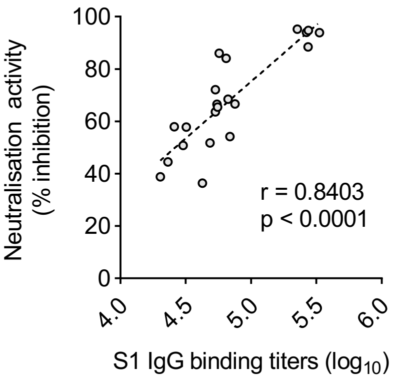


**Supplementary 5: Correlation between S1-specific IgG binding titers and SARS-CoV-2 surrogate neutralising activity.**
C57BL/6J female mice (n=5) were vaccinated IM with $10^7$ PFU of SCV-S in a prime-boost vaccination protocol on days 0 and 28. Serum samples were collected on days 14, 21, and 50 post-vaccination. S1-specific IgG binding titers and neutralization activity were determined by ELISA and cPASS™ SARS-CoV-2 neutralizing antibody detection kit respectively and plotted against each other. A positive correlation was established between the two variables by non-parametric Spearman rank correlation test.

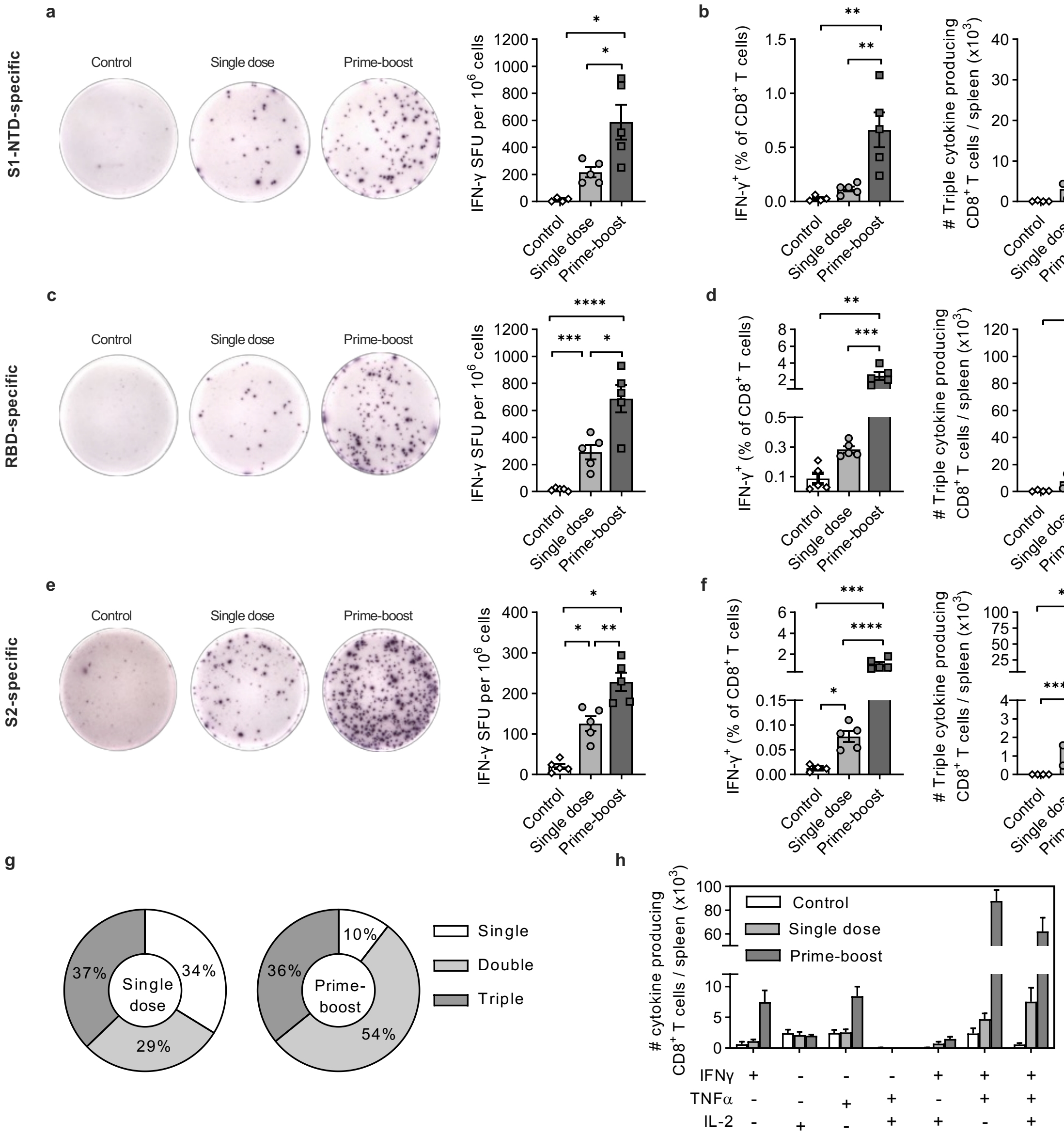


**Supplementary 6: SCV-S induces spike-specific T cell responses in aging mice.** Groups of C57BL/6J aging female mice (9-10 months; n=5) were vaccinated in a single dose (day 0) or prime-boost strategy (day 0 and 28) with SCV-S. Mice vaccinated with control vector on day 0 and 28 were used as controls. At day 120, splenocytes were stimulated with peptide pools (15AA length with 11mer overlaps) spanning the spike protein to examine antigen-specific T cell responses. IFN-γ SFU specific for (**a**) S1 N-terminal domain region, (**c**) RBD region and (**e**) S2 region of the spike protein was quantitated by ELISPOT. Frequency of IFN-$\gamma^+$ $CD8^+$ T cells and absolute numbers of triple cytokine (IFN-$\gamma^+$ $TNF^+$ IL-$2^+$) positive cells for (**b**) S1-NTD region, (**d**) RBD region and (**f**) S2 region of the spike protein was enumerated by intracellular cytokine staining and FACS analysis. Graphs illustrate the (**g**) mean frequency and (**h**) absolute numbers of RBD-specific single, double, and triple cytokine cells within the cytokine positive $CD8^+$ T cell compartment. Symbols represent individual mice and bars indicate mean ± SEM. Data was log transformed and statistical significance determined using Brown-Forsythe and Welch ANOVA with Dunnett T3 multiple comparison test. *$p<0.05$; **$p<0.01$; ***$p<0.001$; ****$p<0.0001$

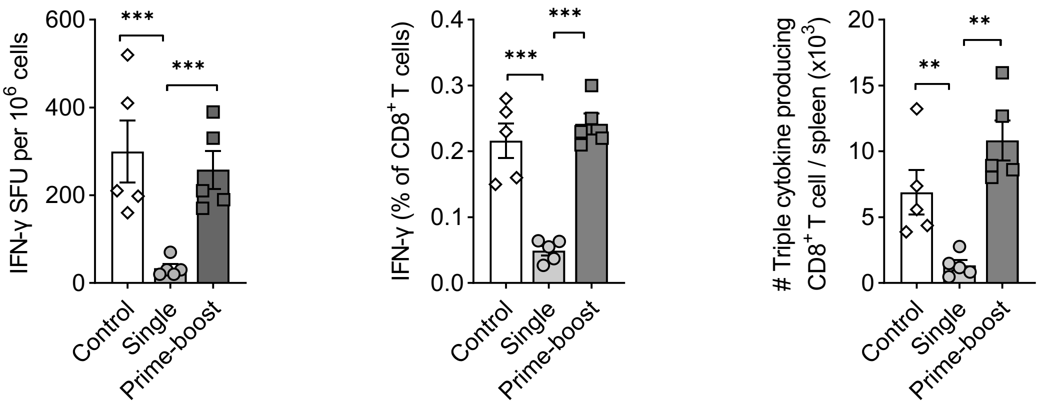


**Supplementary 7: Vector-specific T cell responses increased after boost vaccination.** Groups of C57BL/6J (female; n=5) were vaccinated with SCV-S following a single dose (day 0) or prime-boost strategy (day 0 and 28). Mice vaccinated with control vector on day 0 and 28 were used as antigen-specific controls. At day 120 post-vaccination, splenocytes were stimulated with the VACV-specific A8 peptide (ITYRFYLI) and resultant A8-specific IFN-γ SFUs quantitated by ELISPOT (left panel). Vector-specific $CD3^+$ $CD8^+$ T cell responses were assessed by intracellular cytokine staining and FACS analysis, with the frequency of IFN-$\gamma^+$ $CD8^+$ T cells (middle panel) and absolute numbers of triple cytokine (IFN-$\gamma^+$ $TNF^+$ IL-$2^+$) calculated. Symbols represent individual mice and bars show the mean ± SEM. Data was log transformed and statistical significance was determined using Brown-Forsythe and Welch ANOVA with Dunnett T3 multiple comparison test. *p<0.05; **p<0.01; ***p<0.001; ****p<0.0001

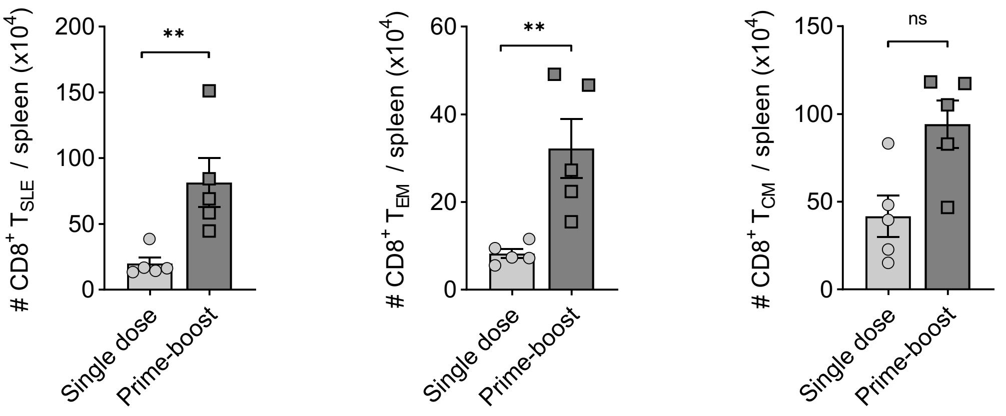


**Supplementary 8: Comparison of effector and central memory CD8 T cell population after single and prime-boost vaccination.**

C57BL/6J female mice (n=5) were vaccinated with SCV-S with a single dose (day 0) or prime-boost regimen (day 0 and 28). At day 120, splenocytes were harvested and the central and effector memory populations were identified by FACS analysis using the surface markers CD3, CD8, KLRG1, CD44, CD62L and CD127. Graphs showing absolute numbers of short-lived effector cells ($T_{SLE}$; $CD44^{hi}$ $KLRG1^{+}$ $CD62L^{lo}$), long-lived effector memory cells ($T^{EM}$; $CD44^{hi}$ $KLRG1^{lo}$ $CD62L^{lo}$ $CD127^{hi}$), and central memory cells ($T_{CM}$; $CD44^{hi}$ $KLRG1^{lo}$ $CD62L^{+}$ $CD127^{hi}$) in single and prime-boost vaccinated mice are shown. Symbols represent individual mice and bars show the mean ± SEM. Data was log transformed and unpaired t test with Welch's correction was used for statistical analysis. **$p<0.01$, ns=not significant

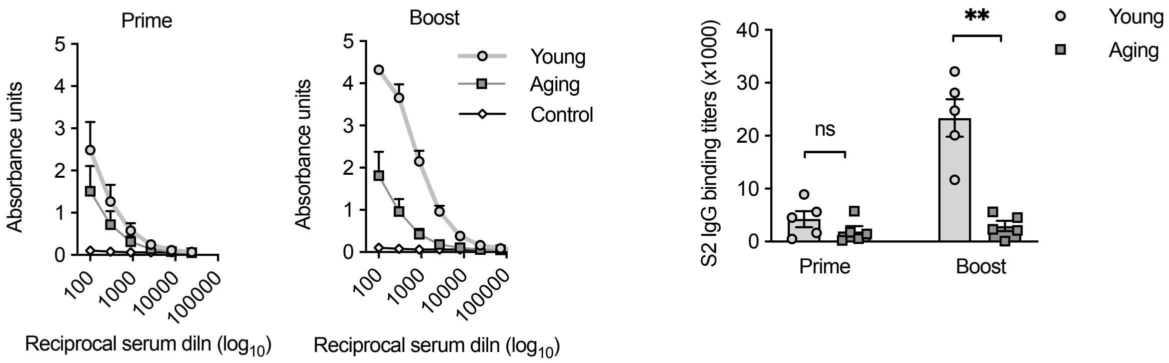


**Supplementary 9: S2-specific binding titers post-vaccination in young and aging mice.** Groups of young (6-8 weeks old) and aging (9-10 months old) female C57BL/6J mice (n=5) were vaccinated IM with $10^7$ PFU of SCV-S (days 0 and 28) or vector control. S2-specific binding activity in serum samples collected on day 21 (prime) and 50 (boost) were evaluated by ELISA. Data was log transformed and statistical significance was determined using unpaired t-test with Welch's correction. **p<0.01

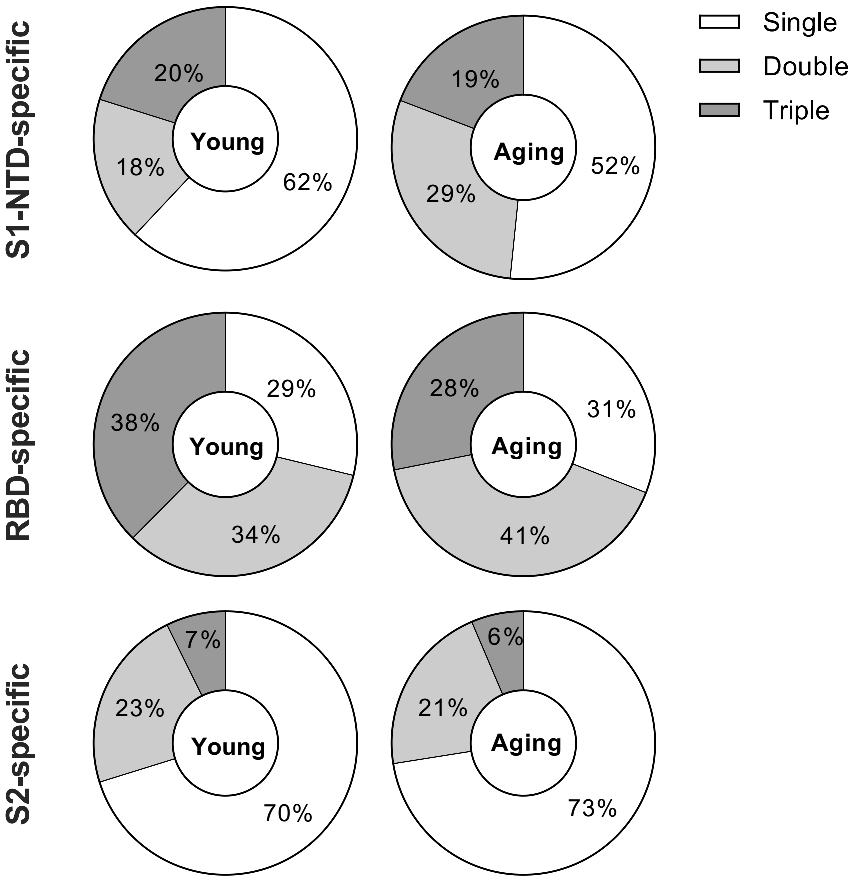


**Supplementary 10: Comparison of multifunctional T cell population in young and aging mice**

Young (6-8 weeks old) and aging (9-10 months old) female C57BL/6J mice (n=5) were vaccinated IM with $10^7$ PFU of SCV-S (days 0 and 28). At 9 months post-vaccination, splenocytes were stimulated with peptide pools (15AA length with 11mer overlaps) spanning the spike protein and the proportions of single and polyfunctional T cells in both the young and aging mice specific to S1 N-terminal domain (top panel), RBD (middle panel) and S2 (bottom panel) were enumerated by intracellular cytokine staining and FACS analysis.
